# Structural and functional characterizations of altered infectivity and immune evasion of SARS-CoV-2 Omicron variant

**DOI:** 10.1101/2021.12.29.474402

**Authors:** Zhen Cui, Pan Liu, Nan Wang, Lei Wang, Kaiyue Fan, Qianhui Zhu, Kang Wang, Ruihong Chen, Rui Feng, Zijing Jia, Minnan Yang, Ge Xu, Boling Zhu, Wangjun Fu, Tianming Chu, Leilei Feng, Yide Wang, Xinran Pei, Peng Yang, Xiaoliang Sunney Xie, Lei Cao, Yunlong Cao, Xiangxi Wang

## Abstract

The SARS-CoV-2 Omicron with increased fitness is spreading rapidly worldwide. Analysis of cryo-EM structures of the Spike (S) from Omicron reveals amino acid substitutions forging new interactions that stably maintain an “active” conformation for receptor recognition. The relatively more compact domain organization confers improved stability and enhances attachment but compromises the efficiency of viral fusion step. Alterations in local conformation, charge and hydrophobic microenvironments underpin the modulation of the epitopes such that they are not recognized by most NTD- and RBD-antibodies, facilitating viral immune escape. Apart from already existing mutations, we have identified three new immune escape sites: 1) Q493R, 2) G446S and 3) S371L/S373P/S375F that confers greater resistance to five of the six classes of RBD-antibodies. Structure of the Omicron S bound with human ACE2, together with analysis of sequence conservation in ACE2 binding region of 25 sarbecovirus members as well as heatmaps of the immunogenic sites and their corresponding mutational frequencies sheds light on conserved and structurally restrained regions that can be used for the development of broad-spectrum vaccines and therapeutics.

## Introduction

Severe Acute Respiratory Coronavirus (SARS-CoV-2) has been continuously evolving through mutations in its viral genome, resulting in increased transmissibility, infectivity and immune escape as observed among emerging variants of concerns (VOCs). Four widely-circulated VOCs, Alpha, Beta, Gamma and Delta, have been previously characterized, among which the Beta variant showed the greatest magnitude of immune evasion from neutralizing antibodies, whereas Delta exhibited dramatically enhanced transmission and infectivity (*1-3*). A newly identified VOC, named the Omicron variant (B.1.1.529), with an unprecedented number of mutations, is quickly spreading worldwide. Although the Delta variant remains the most prevalent strain currently, the Omicron variant is likely to become dominant by early 2022 as predicted by mathematical models (*4*). The Omicron spike (S) harbors over 30 amino acid substitutions, 15 of which are in the receptor binding domain (RBD).

These include three new clusters: 1) S371L, S373P and S375F, 2) N440K and G446S, and 3) Q493, G496, Q498 and Y505H, as well as other accumulated mutations, such as K417N, S477N, T478K, E484A and N501Y (Figure 1A), presumably conferring greater resistance to neutralizing antibodies and vaccine induced humoral immunity (*5-7*). Three small deletions (Δ69-70, Δ143-145, Δ212), one 3-residue insertion (ins214EPE) and four substitutions (A67V, T95I, G142D and N211I) in the N-terminal domain (NTD), alter its local conformation, probably disrupting the original antigenic site existing in the WT and also affecting viral infectivity (Figure 1A). In addition, mutations near by the furin cleavage site like H655Y, N679K and P681H might be implicated in proteolytic activation since the P681R substitution identified in Delta could enhance viral fusogenicity and pathogenicity (*8*). Preliminary data suggests that the Omicron variant escapes almost all clinically approved antibody therapeutics, significantly impairs humoral immunity elicited by natural infection and vaccination, and possesses a higher transmission rates among household contacts than those of the Delta variant, attributing to a higher risk of a yet another resurgence of the pandemic (*9-11*).

**Figure 1.**
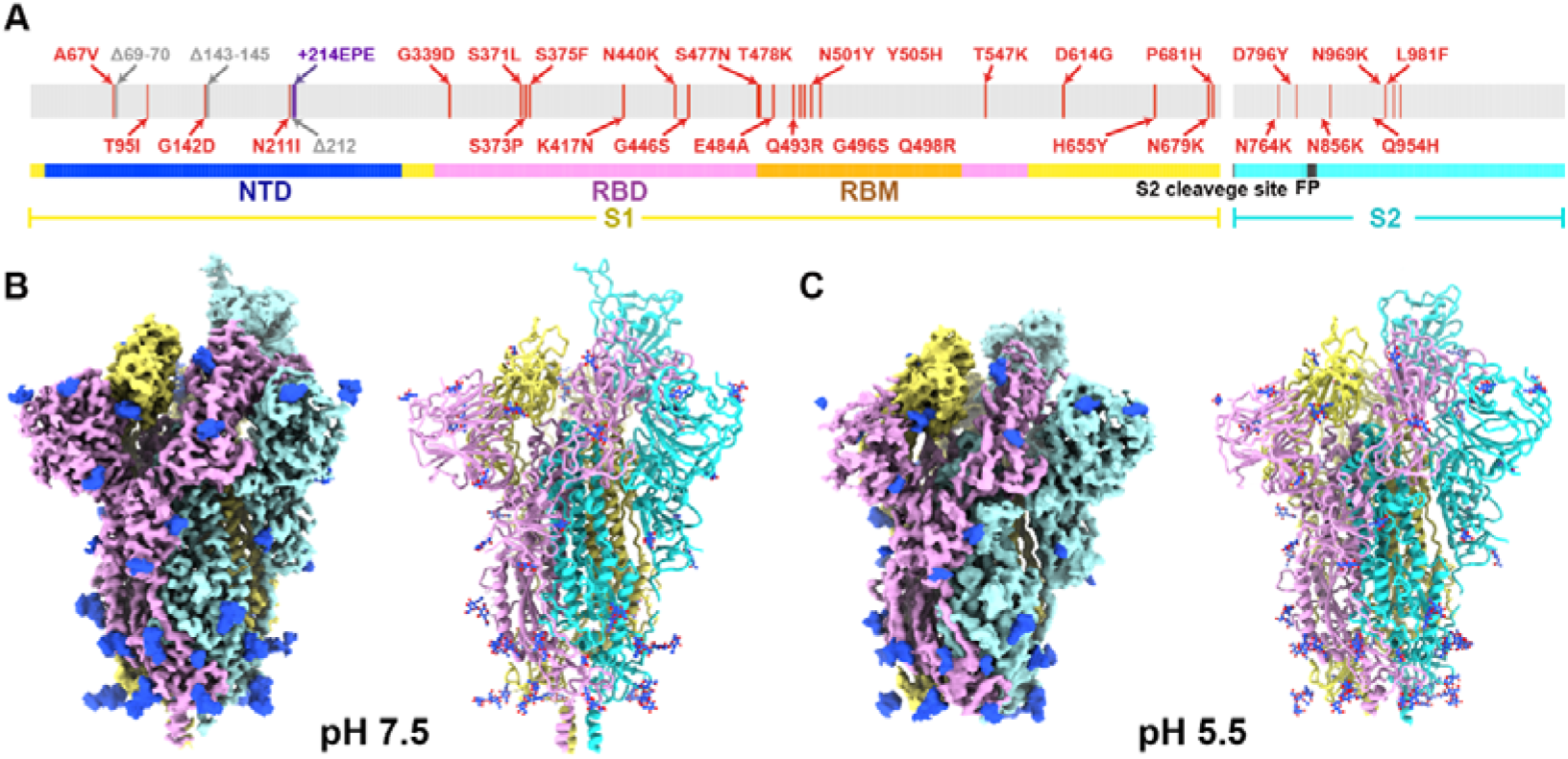
Overall structures of the Omicron S-trimer. (A) Schematic diagram of the full-length SARS-CoV-2 spike (S) protein sequence with substitutions, deletions and insertions are illustrated in red, in grey and purple, respectively. (B and C) Cryo-EM structures of the Omicron S-trimer at serological and endosomal pH. Structures are shown as surface (left) and ribbon (right). The three subunits of S protein are colored in yellow, cyan and magenta, respectively.

SARS-CoV-2 can enter into host cells through endosomes as well as the cellular surface, with inhibition of the activity of both, the endosomal cathepsin L and the cell-surface membrane protein TMPRSS2, required to fully block its entry (*12*). The entry for SARS-CoV-2 involves interactions of S with the ACE2 receptor and known or unidentified attachment cofactors, and subsequent priming of the protein by host cell proteases (*13-15*). These two key events advance the life-cycle of the virus from the prefusion to the postfusion stage, leading to the fusion of the viral membrane with that of the host cell (*12, 16*); details of which are not clearly understood. A set of substitutions, generally comprising of no more than three amino acids, in the RBD of four previous VOCs, have been shown to be associated with an increased affinity towards ACE2(*17*). Surprisingly, the Omicron RBD has nine substitutions (K417N, G446S, S477N, E484A, Q493R, G496S, Q498R, N501Y, Y505H), that are located on the ACE2 binding interface, considerably affecting the receptor recognition, cell tropism and entry pattern, particularly in combination with other mutations, such as H655Y, N679K and P681H (Figure 1A). Thus, understanding the underlying molecular basis for the enhanced transmissibility, immune resistance and virological properties of the Omicron variant may facilitate development of intervention strategies to halt this looming crisis.

## Results

### Cryo-EM structures of the Omicron spike at serological and endosomal pH

Based on the first reported genome sequence of the Omicron variant (Figure 1A), we expressed prefusion-stabilized soluble trimeric ectodomain (S-trimer), which contains GSAS and 6P mutations along with the T4 fibritin trimerization domain (*18, 19*) and purified the protein to homogeneity by affinity chromatography and size-exclusion chromatography (Figure S1). To explore the effect of endosomal pH on triggering conformational alterations, purified S-trimer was dialyzed against PBS buffer (pH 7.5) and acidic solution (0.1M Sodium citrate, pH 5.5), separately. We determined asymmetric cryo-EM reconstructions of the Omicron S-trimer at a resolution of 3.3 Å and 3.9 Å under neutral and acidic pH conditions (Figure 1B, 1C, Figure S1-S3 and Table S1), respectively. Previous cryo-EM structures had revealed two prevalent prefusion conformations (but not limited to two) for SARS-CoV-2 S-trimer, including Alpha, Beta and Kappa variants: a single-up conformation and all-down conformation, related to the positioning of the RBD (*20*). By contrast, only one conformational state with one ‘up’ RBD and two ‘down’ RBDs was observed in structures of the Omicron S-trimer, either at pH 7.5 or at pH 5.5 (Figure 1B and 1C). Distinct from multiple orientations of RBD in the S-trimer at pH 5.5 (*21*), structural heterogeneity in the Omicron S-trimer seems largely decreased, even in endosomal environment. Overall structures of neutral and acidic Omicron S-trimer are indistinguishable, apart from some disorder on the NTD and RBM of the acidic S-trimer (Figure 1B, 1C and Figure S4).

### Structural features underpinning the “up” configuration

To further provide insights into the structural heterogeneity, we also determined near-atomic structures of the Delta S-trimer using the same strategy (Figure S1-S3 and Table S1). In line with WT and most variants, Delta S-trimer exhibits two distinct conformational states corresponding to a closed-form with all three RBDs “down” and an open form with one RBD “up” (Figure S5). To investigate structural basis for the stabilized “up” conformation, we further scrutinized these structures and used the Delta S-trimer for in-depth comparison (Figure 2). Strikingly, a highly compact domain organization at the interface between the “up” S-monomer (defined as mol A) and its adjacent “down” RBD (defined as mol B) was observed in Omicron (Figures 2A and 2B). Compared to the Delta S-trimer, the domain shifts include a ∼15° and ∼8° clockwise rotation of the NTD (mol A, colored in yellow) and “down” RBD (mol B, colored in blue), respectively, together with a counterclockwise rotation of the SD1 (mol A, colored in cyan) and “up” RBD (mol A, colored in violet), underpinning the “up” configuration with buried areas of 450 Å^2^ between two contacted RBDs (Figures 2A and 2B). These extensive contacts were primarily due to hydrophilic and hydrophobic interactions, which were largely mediated by the nine mutations within two patches (Figures 2C-2E). Substitutions of Q493R, G496S, Q498R, N501Y and Y505H in the Omicron RBD (mol B) and K378, Y380, D427 and G413 from the “up” RBD (mol A) established five hydrogen bonds not observed in Delta (Figures 2C and 2D). Furthermore, four-residue mutations with completely opposite characteristics from hydrophilic to hydrophobic, including S371L, S373P, S375F and E484A, together with Y369, A373, F374, V483, G485 and F486 have resulted in massive hydrophobic interactions (Figures 2E and 2F). Previous studies observed that distal regions of the S, such as furin cleavage loop and D614G, can lead to allosteric effects on RBD open/down disposition (*22*). Apart from these, the FPPR segment (fusion peptide proximal region, residues 828-853) and 630 loop (residues 620-640) are also proposed to modulate the S stability and structural rearrangements (*20*). For the Omicron, the FPPR and 630 loop were well solved in the RBD-down conformation, whereas they were partially ordered in the RBD-up conformation (Figure S6), which is also consistent with a recent report that the FPPR segment and 630 loop facilitate clamping down of the RBDs in the closed-form (*20*). These structural observations explain the decreased structural heterogeneity for Omicron S.

**Figure 2.**
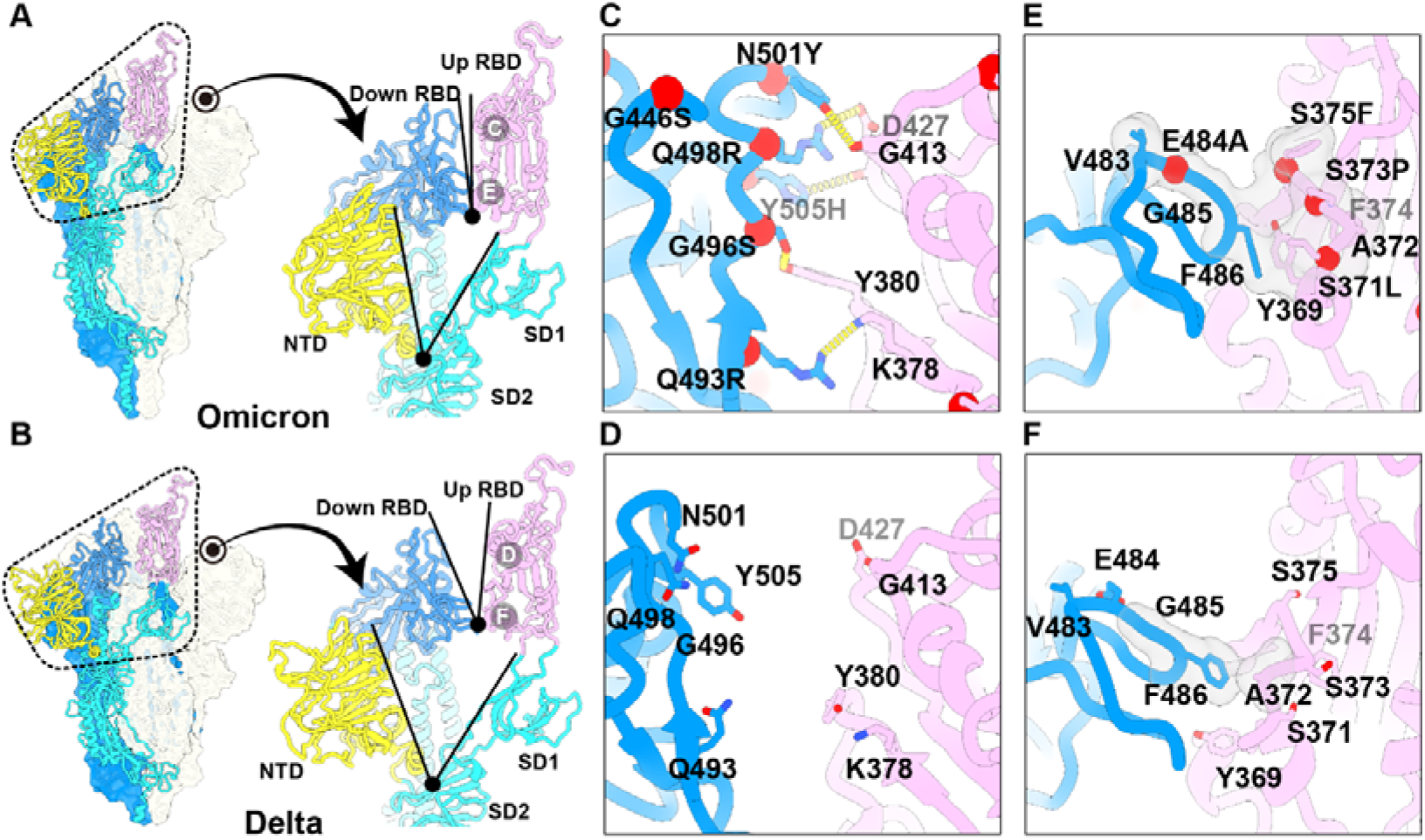
Structural features underpinning the “up” configuration. (A and B) (left) Representation of Omicron (A) and Delta (B) S-trimer in a prefusion conformation with one protomer in ‘open’ state. The ‘up-state’ protomer is shown as ribbon with NTD, RBD and S2 colored in yellow, magenta and cyan, respectively; the neighboring two protomers in ‘down’ state are shown as surface in blue and gray, respectively. (right) Zoomed-in view of the interprotomer RBD-to-RBD contact outlined in black dotted line in (A) and (B) with angles formed by ‘up’ RBD and an adjacent ‘down’ RBD (top), as well as NTD and its inner-protomeric SD1-SD2 axis (bottom) marked out. (C) and (E) Zoomed-in view of interaction details of two independent interfaces for Omicron (A); (D) and (F) zoomed-in view of interaction details of two independent interfaces for Delta (B). The mutated residues are shown as sphere in red with symbols and the residues involved in the interactions are shown as sticks. The hydron bonds are shown as yellow dashed lines (C and D) and hydrophobic network is highlighted in grey (E and F).

### Improved stability and decreased viral fusion ability

Viral stability, particularly surface proteins, like S-trimer, closely correlates with its entry efficiency, viral transmission, adaptability, and immunogenicity (*23*). In spite of adopting an “up” configuration, the Omicron S-trimer exhibits a much more compact architecture in the regions formed by three copies of S2 and RBD, representing a highly stabilized conformation (Figure 3A). Consequently, the Omicron S-trimer possesses substantially increased intersubunit (mol A-mol B) interactions of up to 5,455 Å^2^, when compared to its counterparts from Delta with buried areas of 4,447 Å^2^ (Figure 3B). Similarly, improved intersubunit contacts from either S2-S2 or S1-S1 are also clearly observed, thereby conferring enhanced stability for the Omicron S-trimer in prefusion conformation (Figure 3B). To further investigate the molecular basis for observed tight domain organization in Omicron, superimposition of S2 from Delta and Omicron revealed two local conformational shifts at intersubunit interfaces (residues 849-858 and 968-988) (Figure 3C). In the broader context of the S-trimer, the N856K, N969K and T547K (in SD1) changes from a short side chain to a long one building up three hydrogen bonds with D658 (in SD1), Q755 and S982 from neighboring subunits, pulling the three subunits closer (Figure 3C). Additionally, the substitution of D796Y can stabilize the sugar at residue N709 from its adjacent subunit through forming a hydrogen bond (Figure 3D). In line with structural observations, thermal stability assays verified that the Omicron S-trimer was more stable than those from WT and Delta (Figure 3E).

**Figure 3.**
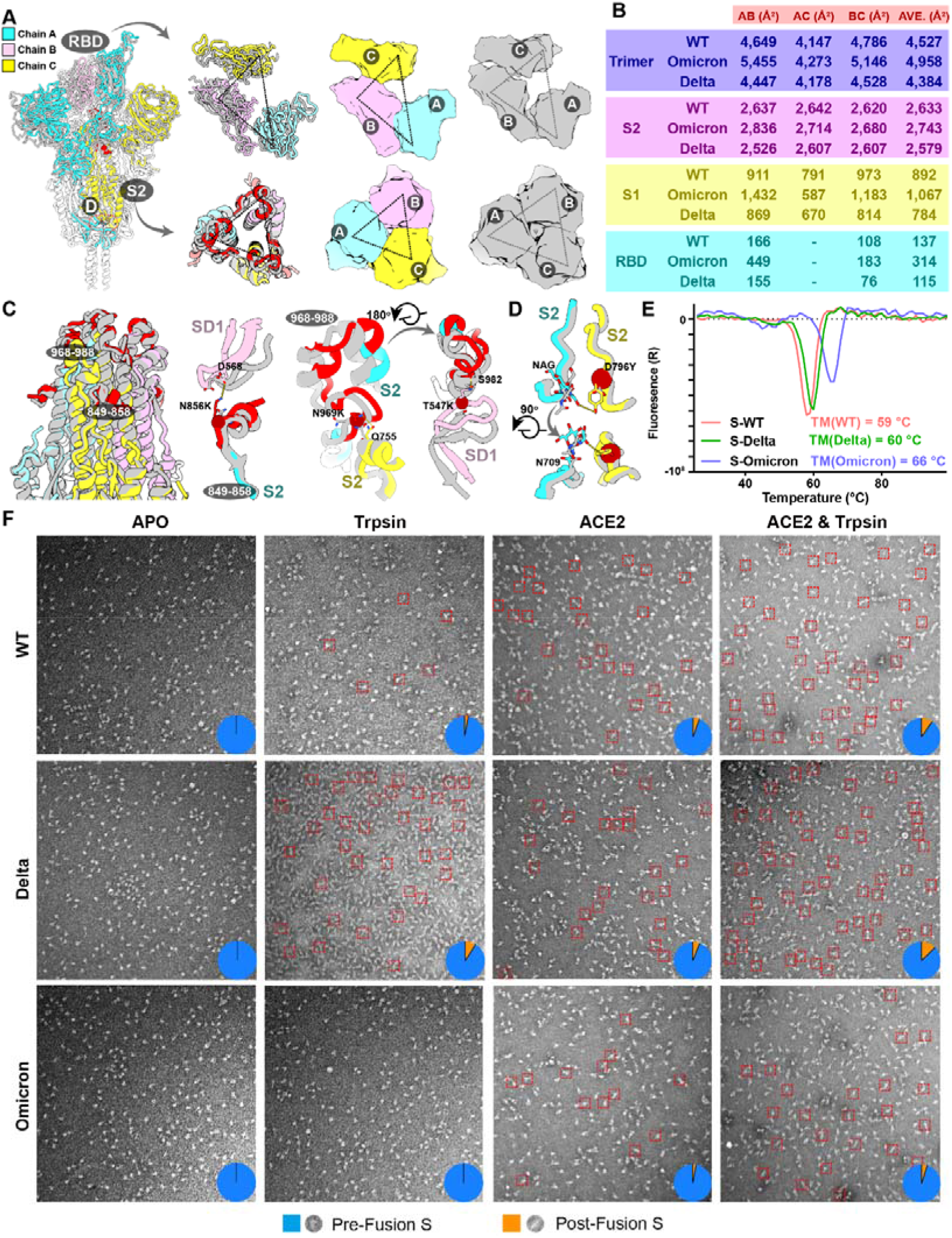
Improved stability and decreased fusogenicity. (A) Superimposition of the structure of the Omicron S-trimer (color scheme is same as in Fig.1B) onto the structure of the Delta S-trimer state (grey). (Right) Top views of the RBD (top) and S2 (bottom) show the inter-subunit contacts of the Omicron and Delta S-trimers. (B) Buried surface areas between two neighboring protomers or S2-, S1-subunits or RBD domains. A, protomer A, B, protomer B, C, protomer C, as labeled in (A). (C) Snapshot of the S2 subunit from (A). Two regions with substantially conformational alterations in Omicron are highlighted in red and shown separately. Residues involved in the formation of hydrogen bonds are shown as sticks. The key mutated residues are shown as sphere in red. (D) The substitution of D796Y in Omicron can stabilize a sugar at N709 from its neighboring protomer. (E) Thermal stability of WT (red), Delta (green) and Omicron (blue) S-trimer. (F) Fusogenicity evaluation of WT, Delta and Omicron S-trimer triggered by treatment of trpsin or trpsin & ACE2. Negative staining images of WT (top), Delta (middle) and Omicron (bottom) S-trimer after incubation with trypsin, ACE2, and both trypsin and ACE2 at conditions described in the method.

In general, improved stability, to some extent, might increase the persistence of the Omicron variant in the exposed environments, posing a higher risk of transmission among household contacts when compared to the Delta variant. Theoretically, improved stability can facilitate viral attachment to the host cells *via* increasing receptor recognition efficiency, however viral membrane fusion may be compromised. To test this, analysis of trypsin- or trypsin & ACE2-mediated S fusogenic conformational rearrangements (*24*) evaluated by negatively stained EM was performed (Figure 3F). Analysis of the negatively stained sample showed that the Omicron S-trimer largely remained in the prefusion conformation and was relatively stable in presence of trypsin & ACE2 at room temperature (Figure 3F). While the same treatment led to formation of more postfusion rosettes for the Delta S-trimer, suggesting that the viral fusion efficiency for the Omicron S-trimer is possibly declined (Figure 3F). These results also largely match the experimental observations of significant reduction in syncitia formation during the Omicron infection (*25*), despite the presence of the P681H substitution, similar change of P681R known to favor S1/S2 cleavage and enhance viral fusogenicity in Delta (*8*).

### Structural basis for altered antigenic characteristics

The overall architectures of the Omicron S-trimer resemble those of WT and other VOCs in the corresponding conformation (Figures 2A and S5). However, the Omicron variant can escape the majority of existing SARS-CoV-2 neutralizing antibodies (NAbs) and humoral immune responses elicited by natural infection or vaccinations, indicative of a considerably altered antigenic structure. The RBD and NTD are two main targets of neutralization in SARS-CoV-2 and other coronaviruses (*26-29*). Superimpositions of RBD and NTD of the WT and Omicron S clearly revealed local conformational alterations in antigenic loops (Figures 4A and 4B). Among these, the most striking differences are in the NTD, which contains three small deletions, one 3-residue insertion and four substitutions (Figure 1A). The mutation A67V together with Δ69-70 triggers the conformational changes in N2 loop. Similarly, N211I, Δ212 as well as ins214EPE alter the configuration of the loop (defined as N4a) encompassing residues 209-216 located adjacent to N4 loop (Figures 4A and S7). Remarkably, the substitution of G142D as well as Δ143-145 leads to a reconfiguration of N3 loop from a hair-pin fold to a loose loop. This reorganization together with the N4a loop, further mediates conformational changes in its neighboring loops N4 and N5 (Figures 4A and S7). Of note, an antigenic supersite comprising N3 and N5 is completely altered in Omicron (*27-29*), structurally explaining the observation that nearly all NTD-targeting NAbs have lost their activities against Omicron. Two major antigenic sites consisting of the 5-6 loop (residues 475-485) and 6-7 loop (residues 492-505) are widely targeted by very potent NAbs via blocking ACE2 binding (Figures 4B and S7), but can be substantially impaired by T478K, E484K and N501Y substitutions observed in several VOCs (*1, 26*). Concernedly, newly occurred mutations, S477N, Q493R, G496S, Q498R and Y505H, together with three already existing mutations, further alter the antigenic characteristics, leading to a striking evasion of the clutches of the antibody. Substitution like S371L, S373P and S375F also cause a clear conformational change in the loop connecting α2 and β2, another antigenic site that is generally conserved across sarbecoviruses (*19, 30*) (Figures 4A and S7). In addition to local conformational alterations, the Omicron RBD and NTD exhibit an increased positively-charged and negatively-charged surface, respectively, when compared to those of WT (Figures 4C-4E).

**Figure 4.**
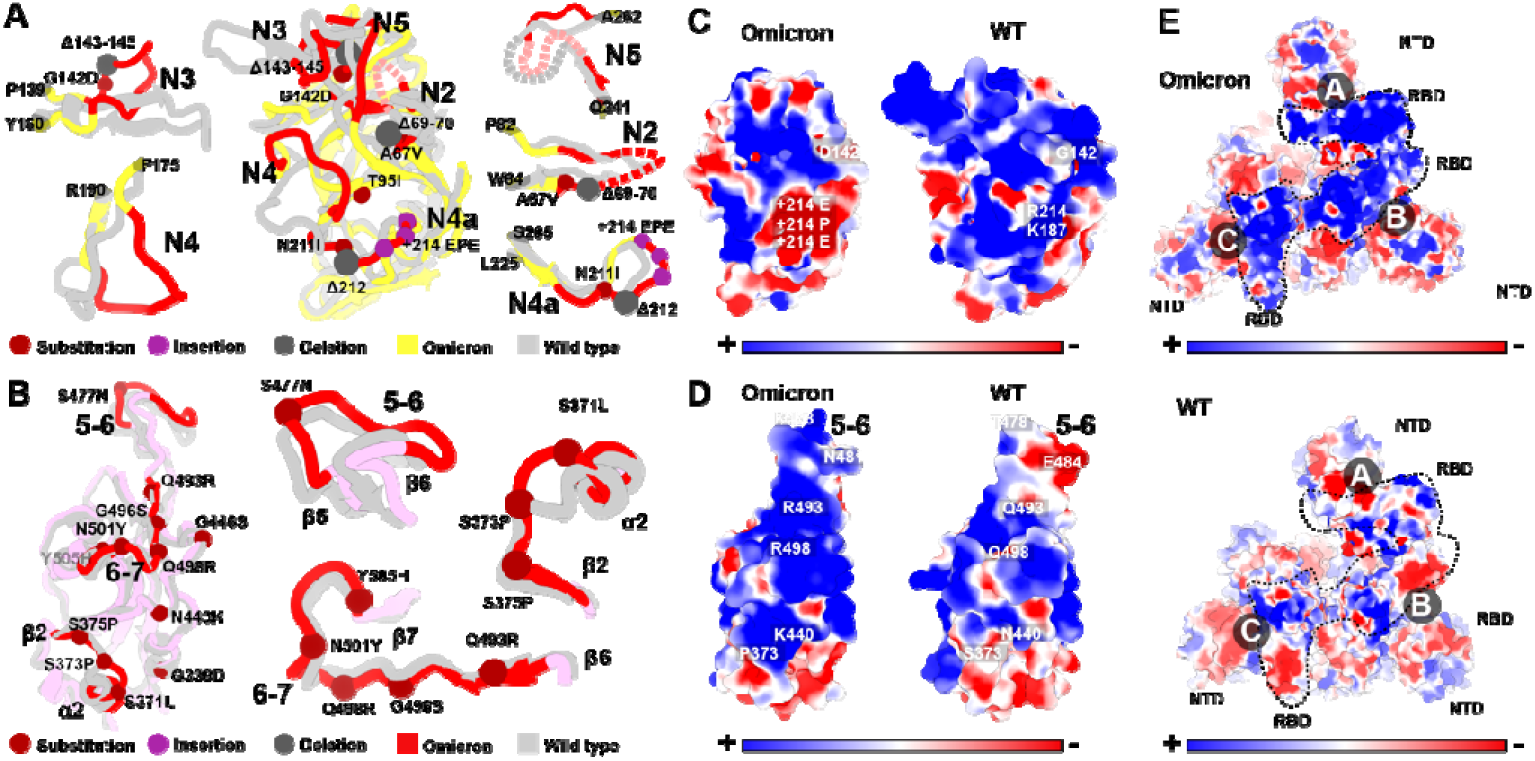
Structural basis for altered antigenic characteristics. (A and B) Superimpositions of the structures of the Omicron NTD (A) and RBD (B) onto WT NTD and RBD. Structures here are shown as ribbon with substitutions, insertions and deletions are depicted as spheres in brown, purple and dark, respectively. Color scheme for Omicron is same as Fig. 1B, the NTD and RBD from WT are colored in grey. Loops of NTD (A) and RBD (B) in Omicron with significantly conformational changes relative to WT were colored in red and zoomed in. (C and D) Electrostatic surface of Omicron (left) and WT (right) NTD (C) and RBD (D). Mutations resulting in dramatically electrostatic changes are labeled. (E) Electrostatic surface of the Omicron (top) and WT (bottom) S-trimer viewed from a top-down perspective. The distinct differences in charge distribution are marked out in dotted lines.

### Structural dissection of the evasion of neutralization of five classes of antibodies

RBD-targeting NAbs can generally be categorized into six classes (from I to VI) based on cluster analysis on epitope from 280 available RBD-NAb complex structures (Figure 5A) (*31*), that are also related to the four/five groups on the basis of competition with the hACE2 for binding to S (*32-34*). We constructed an antigenic heatmap for RBD using the 280 NAb complex structures to estimate *in vivo* antibody recognition frequencies on the RBD (Figure 5B). First three classes of antibodies targeting the RBM with partially overlapped epitopes, are highly potent by way of blocking the interactions between SARS-CoV-2 and ACE2. Class I antibodies, primarily derived from *IGHV3-53/IGHV3-66* with short HCDR3s, recognize only the “up” RBD, and make significant contacts with K417, Q493, N501 and Y505 (Figure 5B). Class III antibodies bound to RBD either in “up” or “down” configuration, extensively associate with E484, Q493 and partially with L452 (Figure 5B). Class II antibodies bind the patch between sites I and III, frequently interacting with S477, T478, E484, Q493 and Y505 (Figure 5B). Class IV antibodies attach to the right shoulder of RBD with relatively condensed epitopes comprising residues 440-450 (Figure 5B). Class V and VI antibodies, generally cross-reactive to sarbecoviruses, target two cryptic epitopes, consisting of residues 351-357, 462-471 and residues 368-385, respectively, which are only accessible when at least one RBD is in the open state (Figure 5B). A number of well-studied NAbs from individual classes were selected to dissect the evasion of neutralization by Omicron (Figure 5C). For Class I NAbs represented by LY-CoV016, substitutions of Q493R and N501Y with longer side chains induced steric clashes with Y102, M101 from HCDR3 and S30 from LCDR1, respectively; mutation K417N further broke the salt bridge with D104 from HCDR3, leading to inactivity in binding to Omicrom S (Figure 5D). Regarding Class II antibodies e.g., REGN10933, changes of K417N and E484A disrupted hydrogen bonds established by D31 from LCDR1 and Y563, S56 from HCDR2, respectively; mutation Q493R also directly clashed with S30 from LCDR1, resulting in the antibody losing its capacity to bind Omicron S (Figure 5D). Similarly, the mutation Q493R caused severe clashes with R104 from HCDR3, and E484A abolished charge interactions with R50 from HCDR2, R96 from LCDR3, resulting in the inability of Class III antibodies exemplified by LY-CoV555 to bind to the Omicron S (Figure 5D). In addition to existing mutations like K417N, E484A and N501Y, Q493R acts as a newly identified key immune escape site, which in conjunction with local conformational changes caused by other nearby mutations, leads to greater resistance to Class I-III antibodies *via* steric clashes. Disruption of the hydrophobic microenvironment constructed by interactions between V445, G447 and P499 from RBD and Y35, V50, Y59 and Y105 from HCDRs of REGN10987, a representative antibody from Class IV antibodies, by the G440S substitution, and the moderate clash between mutation N440K with Y102 from HCDR3, reduces binding affinities of the antibodies to Omicron S dramatically (Figure 5D). As mentioned above, substitutions of S371L, S373P and S375F drove a distinct conformational shift, stabilizing two adjacent RBDs (one up and one down) (Figures 2E and 3B), which consequently altered the relatively conserved antigenic site, mainly targeted by Class VI antibodies (Figure 5D). Fortunately, epitopes of Class V antibodies are mostly beyond the mutated patch, thereby less affected by Omicron, although most NAbs from this class are less potent. In summary, we dissected the evasion of neutralization of the five classes of NAbs by Omicron S and identified three new immune escape sites: 1) Q493R, 2) G446S and 3) S371L/S373P/S375F for antibodies from Classes I-III, IV and VI, respectively.

**Figure 5.**
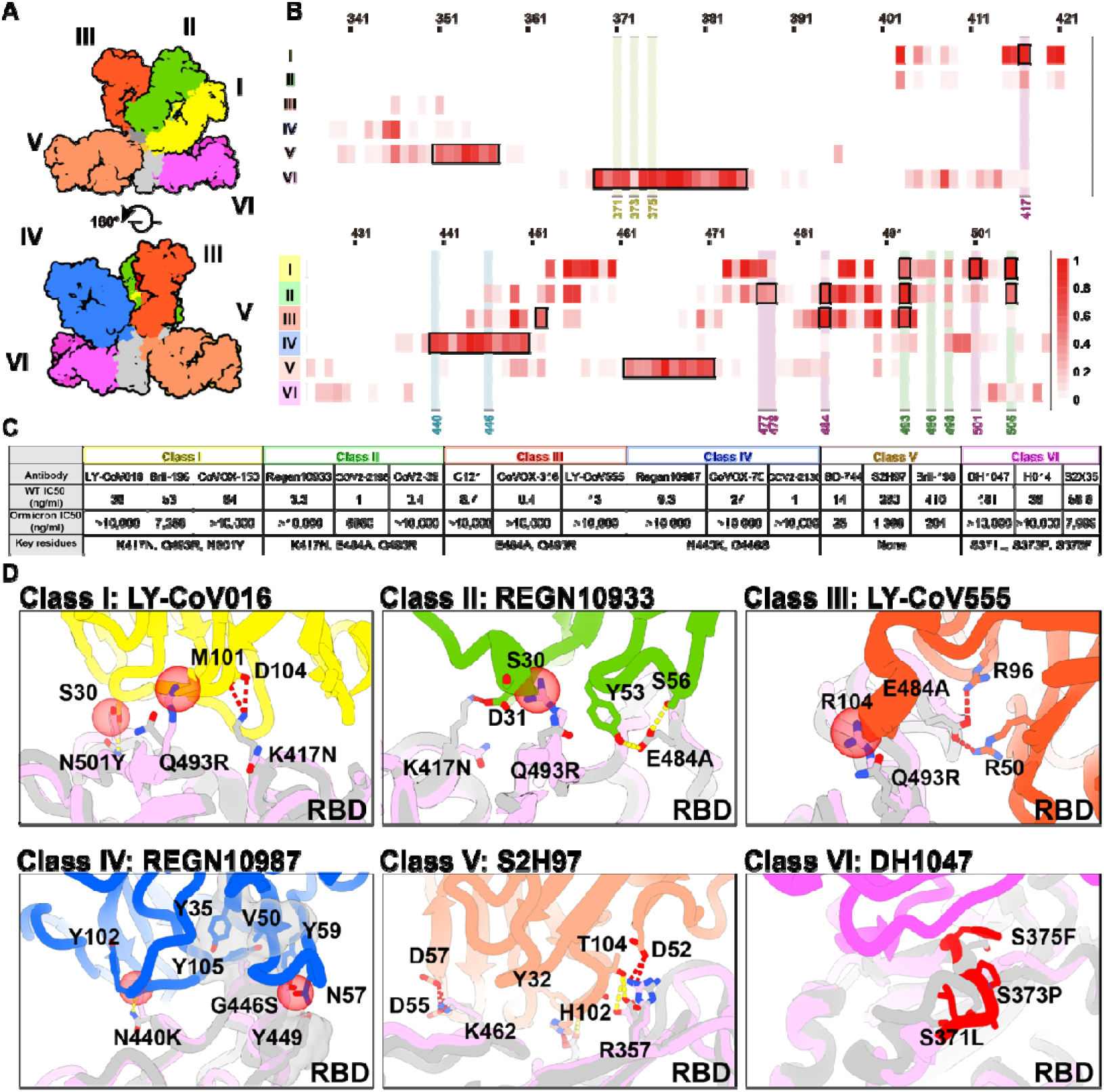
Structural dissection of the evasion of neutralization of antibodies. (A) Surface representation of RBD in complex with six types of NAbs. RBD is colored in grey and the six representative Fab fragments belonging to six classes are colored as follows: class I, yellow; class II, green; class III, red; class IV, blue; V, brown; VI, magenta. (B) Heatmap represents the frequency of RBD residues recognized by NAbs from six classes. Mutations present in Omicron RBD are marked out and highlighted. (C) Summary of representative NAbs from each of six classes. Neutralizing titer (IC_50_) of each NAb against WT and Omicron is enumerated (*5,6,7*). The key residues involved in immune evasion for each class are also listed below. (D) Binding interface between RBD and representative NAbs. All structures are shown as ribbon with the key residues shown with sticks. The clashes between RBD and NAb are shown as red sphere, salt bridges and hydrogen bonds are presented as red dashed lines and yellow dashed lines, respectively. Fab fragments of LY-CoV016, REGN10933, LY-CoV555, REGN10987, S2H97 and DH1047, representatives of Class I, II, III, IV, V and VI, respectively are colored according to the class they belong to; WT RBD is colored in grey; Omicron RBD is colored in light purple.

### Molecular determinants for enhanced binding affinity to human ACE2

Unexpectedly, nine out of fifteen substitutions in the Omicron RBD are located on the human ACE2 binding interface, which considerably affects the receptor recognition. We attempted to measure the binding affinities of WT and Omicron RBD to ACE2 by surface plasmon resonance (SPR) and biolayer interferometry affinities (BLI) (Figure 6A). After the Omicron RBD was loaded onto the biosensor, the human ACE2 (or orthologs of other species) containing buffer solution was passed over the bound RBD. Repeated experiments yielded substantially lower affinities, some even returning negative results, whereas the WT RBD seemed to yield expected binding affinities with ACE2, indicative of the unsuitability of the use of Omicron RBD as immobilization phase. In addition, nonspecific binding of the Omicron RBD to AR2G biosensor was clearly detected when the amine-coupling ACE2 was immobilized onto AR2G biosensor in BLI assay, which also led to unreliable results (Figure S8). Thus, we chose SPR assay, in which the CM5 biosensor was labeled with amine-coupling ACE2 and flooded with the Omicron or WT RBD in the flow through. The Omicron RBD showed a 2.8-fold increased binding affinity to human ACE2 compared to the WT RBD (Figure 6A).

**Figure 6.**
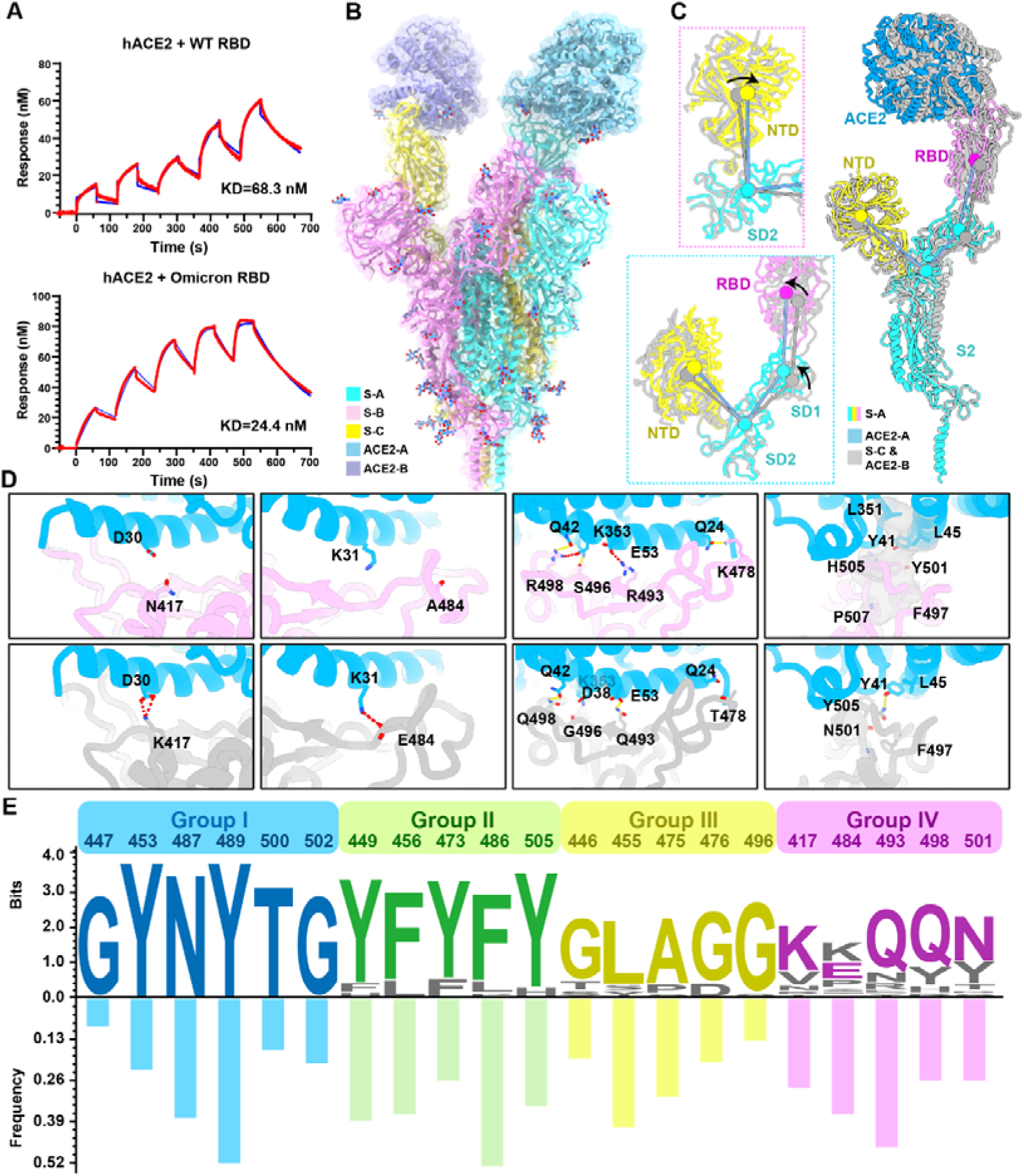
Molecular determinants for enhanced binding affinity to human ACE2. (A) Binding affinity of hACE2 with WT RBD (top) or Omicron RBD (bottom) measured by SPR. (B) Overall structure of the Omicron S-trimer in complex with hACE2. Three copies of S monomer were colored in yellow, cyan and magenta, respectively; two hACE2 molecules bound to RBD were colored in purple and blue, respectively. (C) Superimposition of two S monomer-hACE2 molecules with a focused alignment on S2 subunit. Color scheme for the S monomer in a stabilized “up” conformation is same as Fig. 1B, and the other S monomer is colored in grey. Insets represent the structural shifts of NTD and SD2 (left-top), and altered angles (left-bottom) formed by NTD, SD2, SD1 and RBD, that was triggered by ACE2 binding. (D) Changes at the interfaces between WT RBD and Omicron RBD with hACE2. hACE2, WT RBD and Omicron RBD are colored in blue, grey and magenta, respectively; key mutated residues were shown as sticks with the abolished (left four) and newly-established (right four) bonds denoted in dashed lines; salt bridges, red; hydron bonds, yellow. Hydrophobic network is highlighted in grey. (E) Analysis of sequence conservation and antigenicity frequency on residues involved in ACE2 binding. The logo plot upper represents the conservation of these residues from 25 sarbecoviruses and the histogram below shows the antigenic frequency of the same residues targeted by NAbs. Logos and bars of four types of residues are colored with blue (identical residues), green (homologous residues), yellow (conditionally altered residues) and pink (highly diverse residues), respectively. Amino acid sequences of these residues involved in hACE2 binding are from WT SARS-CoV-2, 17 SARS-CoV-2 variants (Alpha, Beta, Gamma, Delta, Lambda, Mu, Delta plus, Omicron, Eta, Lota, Kappa, Theta, Iota, B.1.1.318, B.1.620, C.1.2, C.363), SARS-CoV-1, Pangolin coronavirus (GX/P2V/2017 and GD/1/2019), Bat coronavirus (WIV, RaTG13 and LYRa11) and Civet coronavirus of 007/2004.

To further unveil molecular details of the binding, we determined cryo-EM structures of the Omicron S-trimer in complex with human ACE2 at 3.5 Å (Figures 6B, S1-S3 and Table S1). The structure showed two copies of ACE2 bound to two RBDs (mol A and mol C) in the “up” conformation. The stabilized “up” S-monomer showed no notable conformational alterations upon ACE2 binding, but the other “up” S-monomer had a wider angle between NTD, SD2, SD1 and RBD, that was triggered by ACE2 (Figure 6C), suggesting that the stabilized “up” configuration is tolerant to the interference by ACE2 binding, probably antibody binding as well. Local refinement of the RBD-ACE2 region resulted in a reliable density map for analysis of the mode of interaction between Omicron S and ACE2 (Figures S1-S3). Structural comparisons of the key interactions at the interface between the Omicron/WT and ACE2 revealed that new substitutions of T478K, Q493R, G496S and Q498R strengthened the binding of Omicron to ACE2 by establishing hydrogen bonds or salt bridges with Q24, E35, K353 and D38, respectively (Figure 6D). Together with the N501Y, a mutation known to improve the binding affinity by 6-fold (*17*), these mutations facilitate the binding of Omicron to ACE2. Meanwhile, mutations K417N and E484A decreased binding affinity to ACE2 via breaking two hydrogen bonds in Omicron. This reduction in affinity is offset by new interactions forged by other mutations. Overall the Omicron variant possesses improved binding to ACE2.

## Discussion

Viruses latch on to receptors using conserved residues. Many members of the β-Coronavirus lineage B (termed as sarbecoviruses), including SARS-CoV-1, SARS-CoV-2 variants, civet, bat and pangolin derived sarbecoviruses, can utilize human ACE2 to enter host cells (*35, 36*). Interestingly, SARS-CoV-2 variants and sarbecoviruses from pangolin have higher affinities for ACE2 than SARS-CoV-1, and sarbecoviruses originating from civets as well as bats, which might be explained by the affinity-enhancing mutations present in these viruses (*36, 37*). An analysis of the conservation of protein sequence around the receptor binding sites of 25 reported sarbecovirus members relying on human ACE2 for cellular entry reveals that 11 residues of the virus out of a total of 21 residues that engage with the receptor are highly conserved (Figure 6E). The amino acids of the virus involving in binding ACE2 can be categorized into four groups based on their conservation and essential roles in binding: 1) Group I includes six identical residues such as G447, Y453, N487, Y489, T500 and G502, 2) Group II consists of five homologous residues like Y449/F/H, F456/L, Y473/F, F486/L and Y505H, 3) Group III has five conditionally altered residues like G446/S/T, L455/S/Y, A475/P/S, G476/D and G496/S, 4) Lastly, Group IV includes five highly diverse residues K417/V/N/R/T, E484/K/P/Q/V/A, Q493/N/E/R/Y, Q498/Y/H/R and N501/Y/T/D/S (Figure 6E). Mutations in the first two groups of amino acids are strictly constrained, presumably because they act as molecular determinants in either retaining basic affinity for ACE2 or ensuring proper protein folding, while substitutions in the last two groups of amino acids are, to some extent, tolerated or even enhance ACE2 binding, and are frequently observed in various VOCs (Figure 6E). Of note, amino acids belonging to the Group IV that dramatically affect affinity include residues at position 417 (e.g., K417), 484 (e.g., K484), 493 (e.g., R493), 498 (e.g., R498) and 501 (e.g., Y501). These amino acids are also known to vary across sarbecoviruses. Furthermore, consensus mutations, including G446T/S, F456L, Y473F and G476D are observed in low affinity group (SARS-CoV-1, civet and bat sarbecoviruses), indicative of a concerted role for these mutations in regulating affinity for ACE2 (Figure S9). In addition, the sequence and structural analysis revealed substantial plasticity in a complicated network formed by multiple substitutions, as some mutations probably increase polar contacts, while others may impair hydrophobic interactions. Our results are largely supported by deep mutational scanning assays, unveiling the details of molecular interactions involved in ACE2 binding (Figure S10) (*37*).

In addition to the pursuit of greater transmissibility and infectivity, viral evolution is primarily driven by immune escape as well. Our immunogenic and mutational heatmaps for RBD using the 280 NAb complex structures to estimate *in vivo* NAb-targeting frequencies and viral mutation frequencies revealed an overall positive correlation between “hot” immunogenic sites and areas with high mutation frequencies, but with an exception for some sites involved in ACE2 binding (Figure 6E). Only 3 of the top 10 hottest immunogenic residues contained substitutions, (Q493R, E484K/A, Y505H) in circulating SARS-CoV-2 variants, among which Q493R and Y505H are newly acquired in Omicron (Figure 6E). Particularly, “hot” immunogenic residues, such as F486, Y489, Y449, N487 and F456, are mostly from the Groups I and II type of residues within ACE2 binding sites that have never undergone mutagenesis during selection, even in presence of maximal immune pressure exerted during the ongoing COVID-19 pandemic (Figure 6E). In other words, substitutions of these “hot” immunogenic sites are best avoided because they probably sacrifice binding activity to ACE2, which could prove to be fatal to the virus. These correlates clearly explain why mutations at these positions would not be selected. In line with our findings, several ACE2-mimic antibodies have already been shown to broadly cross-neutralize sarbecoviruses (*7, 38*). The conserved and structurally constrained region for ACE2 recognition revealed in this study would rationalize the development of broad-spectrum vaccine and antibody therapeutics.

## Materials and Methods

### Cell lines

HEK293T cells (ATCC, CRL-3216) were cultured in Dulbecco’s Modified Eagle’s Medium (DMEM) supplemented with 10% fetal bovine serum (FBS). The cultures were maintained at 37 °C in an incubator supplied with 8% CO2.

### Protein expression and purification

The plasmids encoding the full-length spike (S) protein (residues 1-1028) of wild-type SARS-CoV-2 (GenBank: MN908947) was used as template for construction of the S gene of Delta (T19R, G142D, EF156-157del, R158G, L452R, T478K, D614G, P681R, D950N) and Omicron (A67V, Δ69-70, T95I, G142D, Δ143-145, Δ211, L212I, ins214EPE, G339D, S371L, S373P, S375F, K417N, N440K, G446S, S477N, T478K, E484A, Q493K, G496S, Q498R, N501Y, Y505H, T547K, D614G, H655Y, N679K, P681H, N764K, D796Y, N856K, Q954H, N969K, L981F) by overlapping PCR. All the full-length S gene constructs have six proline substitutions at residues 817, 892, 899, 942, 986 and 987 and two alanine substitutions at residues 683 and 685 and a C-terminal T4 fibritin foldon domain to facilitate the protein expression and stabilization of the trimer conformation (ACROBiosystems, Cat No. SPN-C52Hz). All the constructs described above were attached with a C-terminal six-His for protein purification. To obtain these proteins, the plasmids constructed above were transiently transfected into HEK293 F cells grown in suspension at 37°C in a rotating, humidified incubator supplied with 8% CO2 and maintained at 130 rpm. After incubation for 72 hours, the supernatant was harvested, concentrated and exchanged into the binding buffer by tangential flow filtration cassette. The protein of interest was separated by affinity chromatography using resin attached with Ni-NTA and being subjected to additional purification by size exclusion chromatography using a Superose 6 10/300 column (GE Healthcare) in 20 mM Tris pH 8.0 and 200 mM NaCl.

### S trimer thermal stability release assay

PaSTRy was performed with SYPRO Orange (Invitrogen, Carlsbad, USA) as fluorescent probes to detect the exposed hydrophobic residues by an MX3005 qPCR instrument (Agilent, Santa Clara, USA). Here, we set up pH=7.4 and pH=4.5 25 μl reaction system which contained 10 μg of target protein i.e., S trimer of Wild-type, Delta and Omicron, 1000x SYPRO Orange, and ramped up the temperature from 25 °C to 99 °C. Fluorescence was recorded in triplicate at an interval of 1 °C.

### Bio-layer interferometry

Bio-layer interferometry (BLI) experiments were run on an Octet Red 96e machine (Fortebio). To measure the binding affinities of RBD from different variants with ACE2, His-tagged wild-type RBD, Delta RBD or Omicron RBD were immobilized onto NTA biosensors (Fortebio) and threefold serial dilutions of ACE2 were used as analytes. Also, the ACE2 was immobilized onto AR2G biosensors (Fortebio), as RBD of wild-type, Delta and Omicron as the analytes. Data were then recorded using software Data Acquisition 11.1 (Fortebio) and analyzed using software Data Analysis HT 11.1 (Fortebio) with a 1: 1 fitting model.

### Surface plasmon resonance

His-tagged Wild-type S, Delta S, Omicron S, WT RBD, Delta RBD and Omicron RBD were immobilized onto Ni-NTA sensor chip (GE Healthcare) using a Biacore X100 (GE Healthcare). Serial dilutions of purified ACE2 were injected ranging in concentrations from 250 to 15.6 nM. The resulting data were fitted to a 1:1 binding model using Biacore Evaluation Software (GE Healthcare). Also the ACE2 was immobilized onto CM5 chip and Serial dilutions of purified WT, Delta or Omicron RBD were injected ranging in concentrations from 250 to 15.6 nM.

### Conformational change analysis using negative staining EM

The WT, Delta and Omicron S-trimer at 1 mg/ml were incubated with four times the molar of hACE2 overnight on ice. Samples were diluted to a concentration of 0.03 mg/ml before being imaged and negatively stained for EM. For the treatment of trypsin, samples were added with 1.6 μg/ml trypsin at room temperature for 15 min and used for imaging by negative staining EM.

### Cryo-EM sample preparation, data collection

The Delta S-trimer, Omicron S-trimer (under neutral or acidic pH) or Omicron S trimer (under neutral pH) mixed with hACE2 was dropped onto the pre-glow-discharged holey carbon-coated gold grid (C-flat, 300-mesh, 1.2/1.3, Protochips In.), blotted for 6 seconds with no force in 100% relative humidity and immediately plunged into the liquid ethane using Vitrobot (FEI). Cryo-EM data sets of these complexes were collected at 300 kV with an FEI Titan Krios microscope (FEI). Movies (32 frames, each 0.2 s, total dose of 60 e− A□-2) were recorded using a K3 Summit direct detector with a defocus range between 1.5-2.5 μm. Automated single particle data acquisition was carried out by SerialEM, with a calibrated magnification of 22,500 yielding a final pixel size of 1.07 A□.

### Cryo-EM data processing

A total of 3,106, 1,777, 6,050 and 5,030 micrographs of Delta S-trimer, Omicron S-trimer under neutral and acidic pH conditions and neutral Omicron S-trimer mixed with hACE2 were recorded. All the micrographs were processed with MotionCor2 in Relion3.0. The CTF value of each micrograph was estimated by Gctf. Then 1,317,048, 830,943, 1,789,492 and 916,297 particles of Delta S-trimer, Omicron S-trimer under neutral and acidic pH conditions and neutral Omicron S-trimer mixed with hACE2 were picked and extracted by Relion3.0. Apo S and S-trimer-hACE2 complex were extracted with a 320^2^-pixel box and a 360^2^-pixel box, respectively. Reference free 2D alignment by cryoSPARC (*39*) was applied for all the particles. Based on the results of 2D alignment, 926,504 and 458,926 particles of Delta S-trimer and neutral Omicron S-trimer complexed with hACE2 were selected and applied for Heterogeneous Refinement by cryoSPARC. Then 309,137 and 479,164 particles of Omicron S-trimer under neutral and acidic pH were selected and applied for template-guided 3D-classification by Relion3.0 (*40*). For all the classifications, no symmetry was imposed. When the potential conformation for the each structure was produced, particles from each candidate model were selected and processed by non-uniform auto-refinement and postprocessing in cryoSPARC to generate the final cryo-EM density for Delta S-trimer, Omicron S-trimer under neutral and acidic pH conditions and neutral Omicron S-trimer-hACE2 complex. To improve the resolution of the interface between RBD and hACE2, the local refinement (*41, 42*) was used to obtain the final resolution of the focused interfaces which contained the interfaces of RBD and hACE2 investigated here as described previously. The resolution was determined based on the gold-standard Fourier shell correlation (threshold = 0.143) and evaluated by ResMap (*43*). All dataset processing is shown in Fig.S2 and also summarized in Table 1.

### Model fitting and refinement

The atom models of the complexes were generated by first fitting the chains of native apo SARS-CoV-2 S trimer (PDB number of 6VYB) and ACE2 (PDB number of 6M0J) into the obtained cryo-EM densities by Chimera. Then the structure was manually adjusted and corrected according to the protein sequences and cryo-EM densities in Coot and finally real-space refinement was performing by Phenix. Details of the refinement statistics of the complexes are summarized in Table 1.

### Data availability

The atomic coordinates of neutral Omicron variants Spike trimer, acidic Omicron variant Spike trimer, Delta variant Spike trimer (3RBD down), Delta variant Spike trimer (1 RBD up) and neutral Omicron variant Spike trimer in complex with human ACE2 have been submitted to the Protein Data Bank with accession numbers:7WG6, 7WG7, 7WG8, 7WG9 and 7WGB, respectively. Cryo-EM density maps have been deposited at the Electron Microscopy Data Bank with accession codes EMD-32478, EMD-32479, EMD-32480, EMD-32481 and EMD-32482. To reveal structural details of interactions between Omicron variant Spike trimer and human ACE2, the local optimized reconstruction is used, and the related atomic model and EM density map are deposited under accession code 7WGC and EMD-32483, respectively.

## Acknowledges

We thank Dr. Xiaojun Huang, Dr. Xujing Li and Dr. Lihong Chen for cryo-EM data collection at the Center for Biological imaging (CBI) in Institution of Biophysics, CAS. We also thank Dr. Yuanyuan Chen, Zhenwei Yang, Bingxue Zhou for technical support on BLI and SPR. Work was supported by the Strategic Priority Research Program (XDB29010000, XDB37030000), CAS (YSBR-010), National Key Research and Development Program (2020YFA0707500, 2018YFA0900801), Beijing Municipal Science and Technology Project (Z201100005420017) and Ministry of Science and Technology of China (EKPG21-09 and CPL-1233). Xiangxi Wang was supported by Ten Thousand Talent Program and the NSFS Innovative Research Group (No. 81921005). Kang Wang was supported by the Special Research Assistant Project of the Chinese Academy of Medical Sciences.

## Author contributions

X.W., Y.C., and L.C. designed the whole study; Z.C., P.L., N.W., L.W., K.F., Q.Z., K.W., R.C., R.F., Z.J., M.Y., G.X., B.Z., W.F., T.C., L.F., Y.W., X.R., P.Y performed experiments; P.L., N.W., L.W., L.C. prepared the Cryo-EM samples and solved the structures; all authors analyzed data; X.W., Y.C., and L.C wrote the manuscript with input from all co-authors.

## Supplemental figures, titles and legends

**Figure S1.**
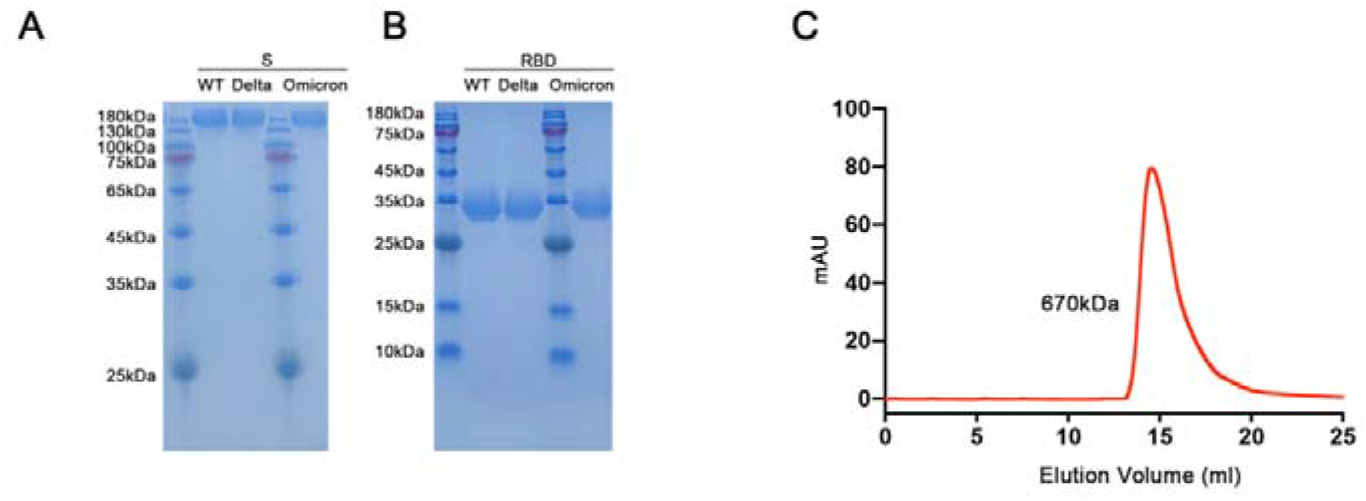
Purification and characterization of Wild-type, Delta and Omicron S trimer and RBD. (**A**) SDS-PAGE analysis of the Wild-type, Delta and Omicron S trimer. (**B**) SDS-PAGE analysis of the Wild-type, Delta and Omicron RBD. (**C**) Gel filtration profile of the affinity-purified Omicron S trimer.

**Figure S2.**
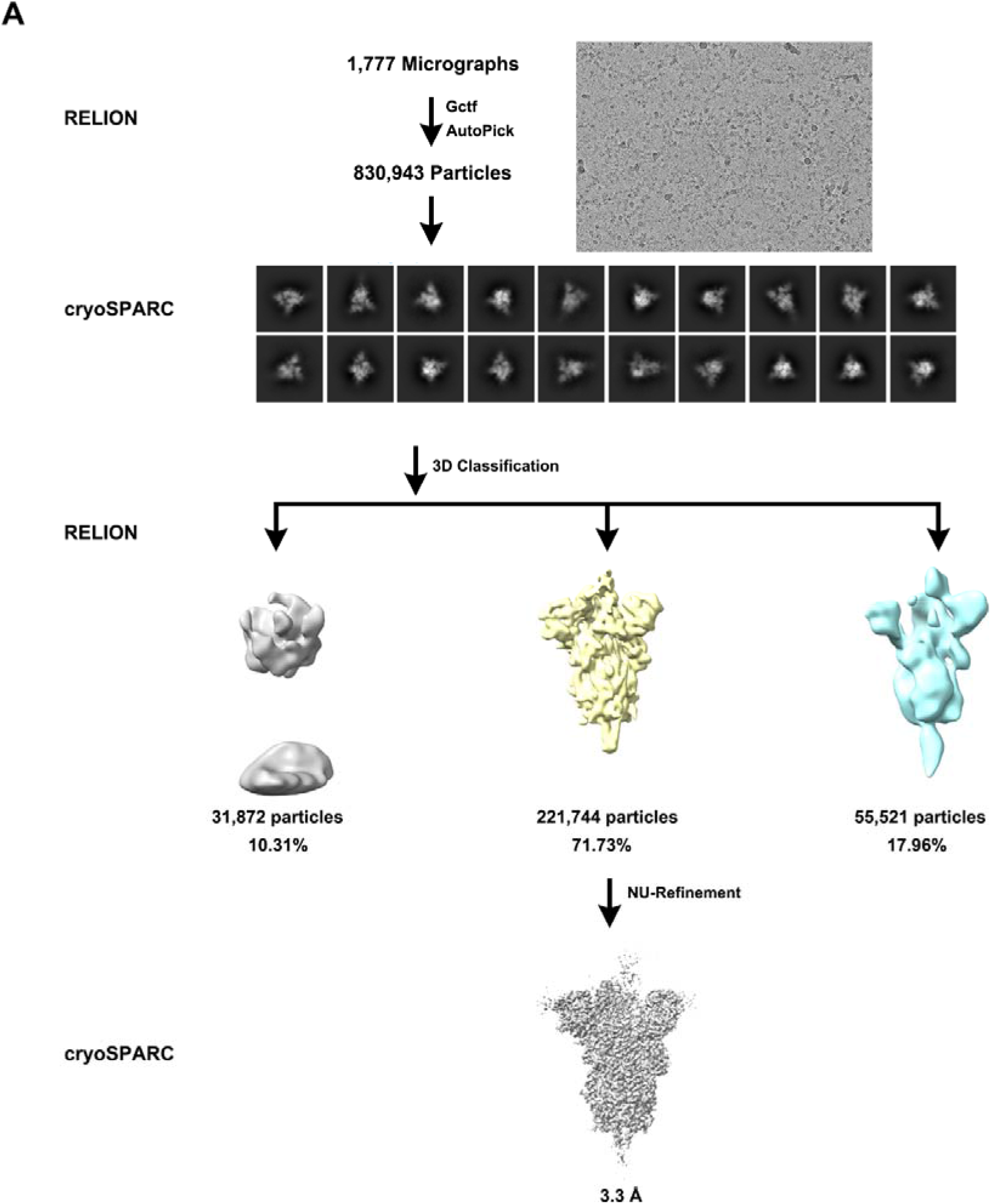
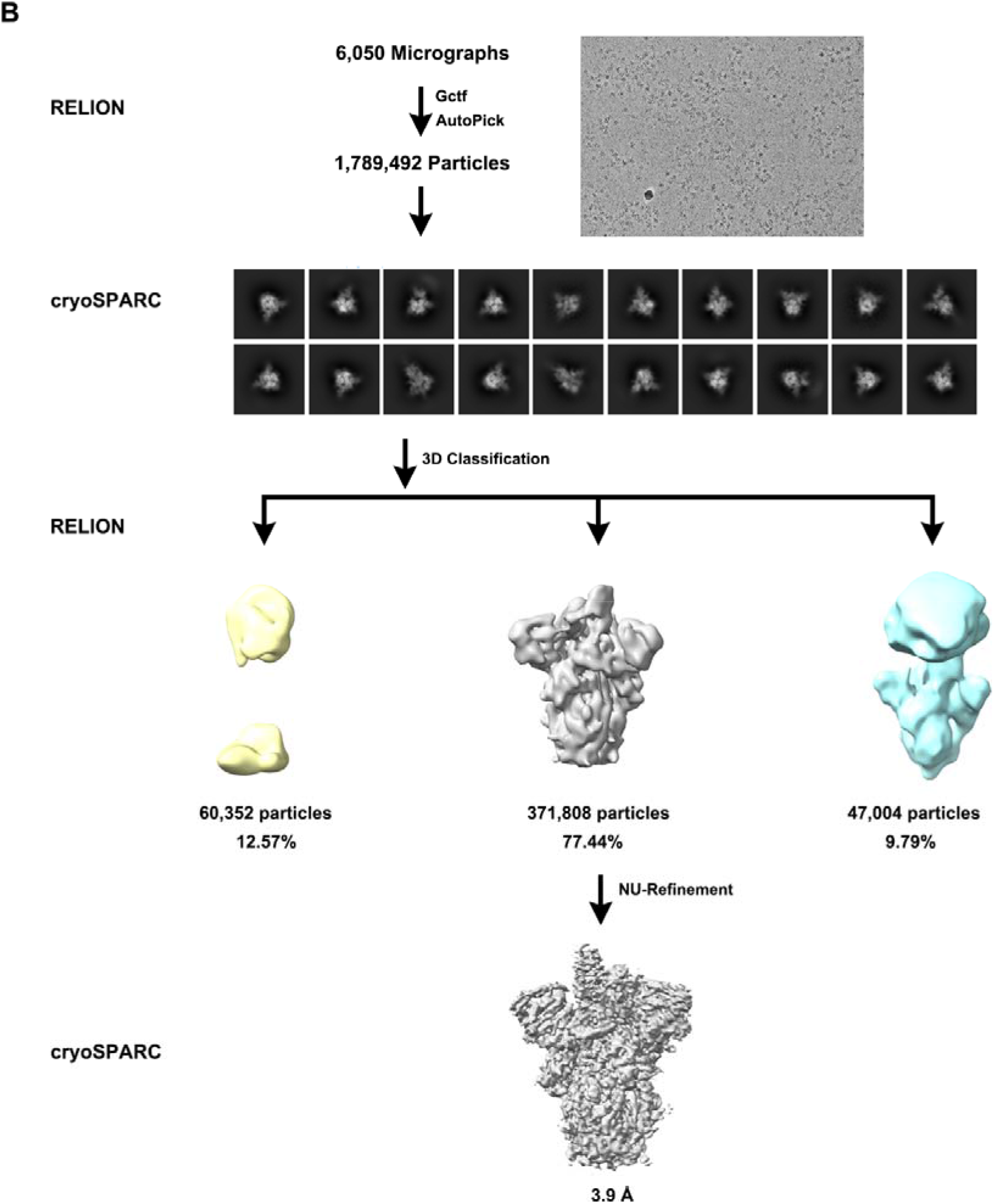
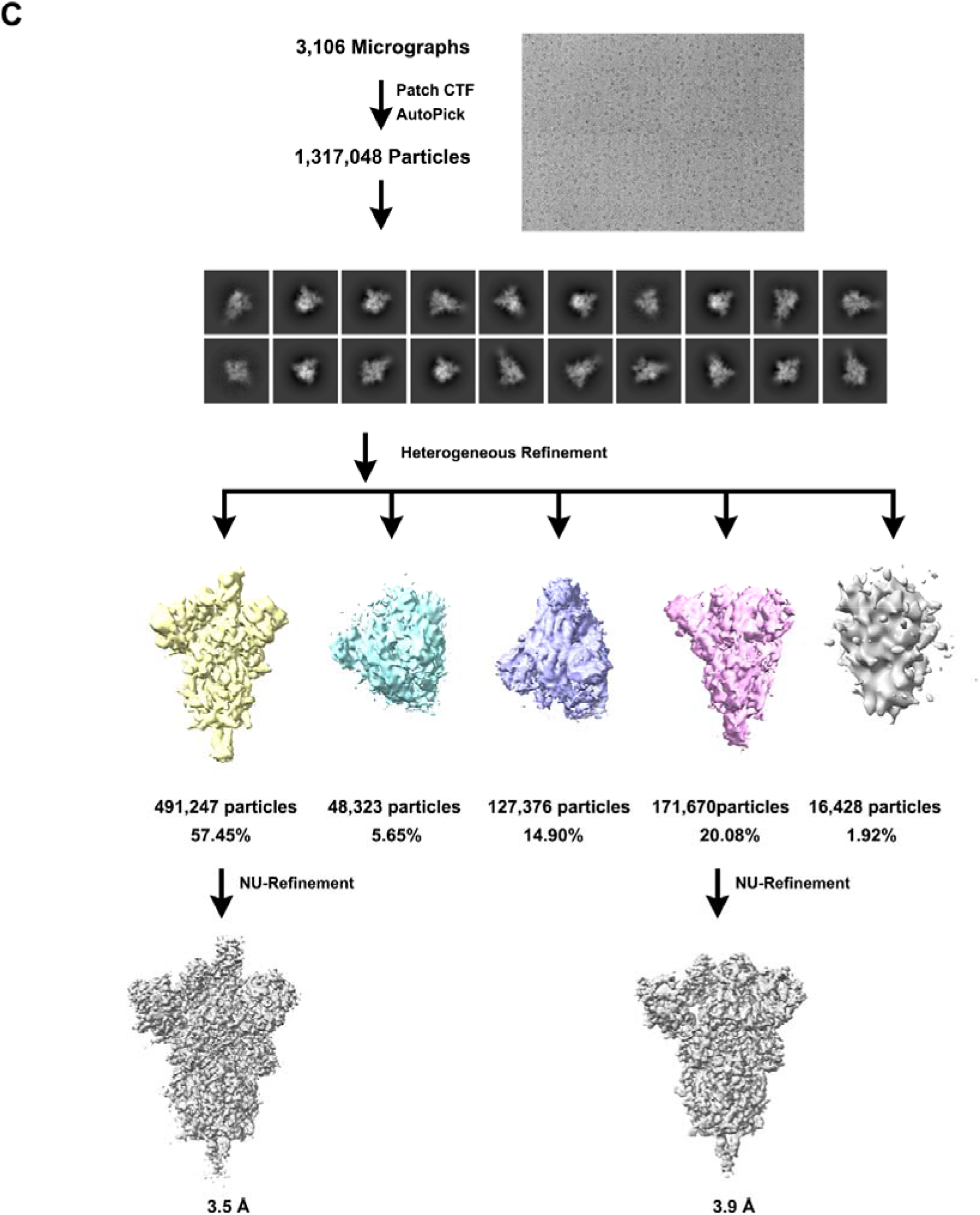
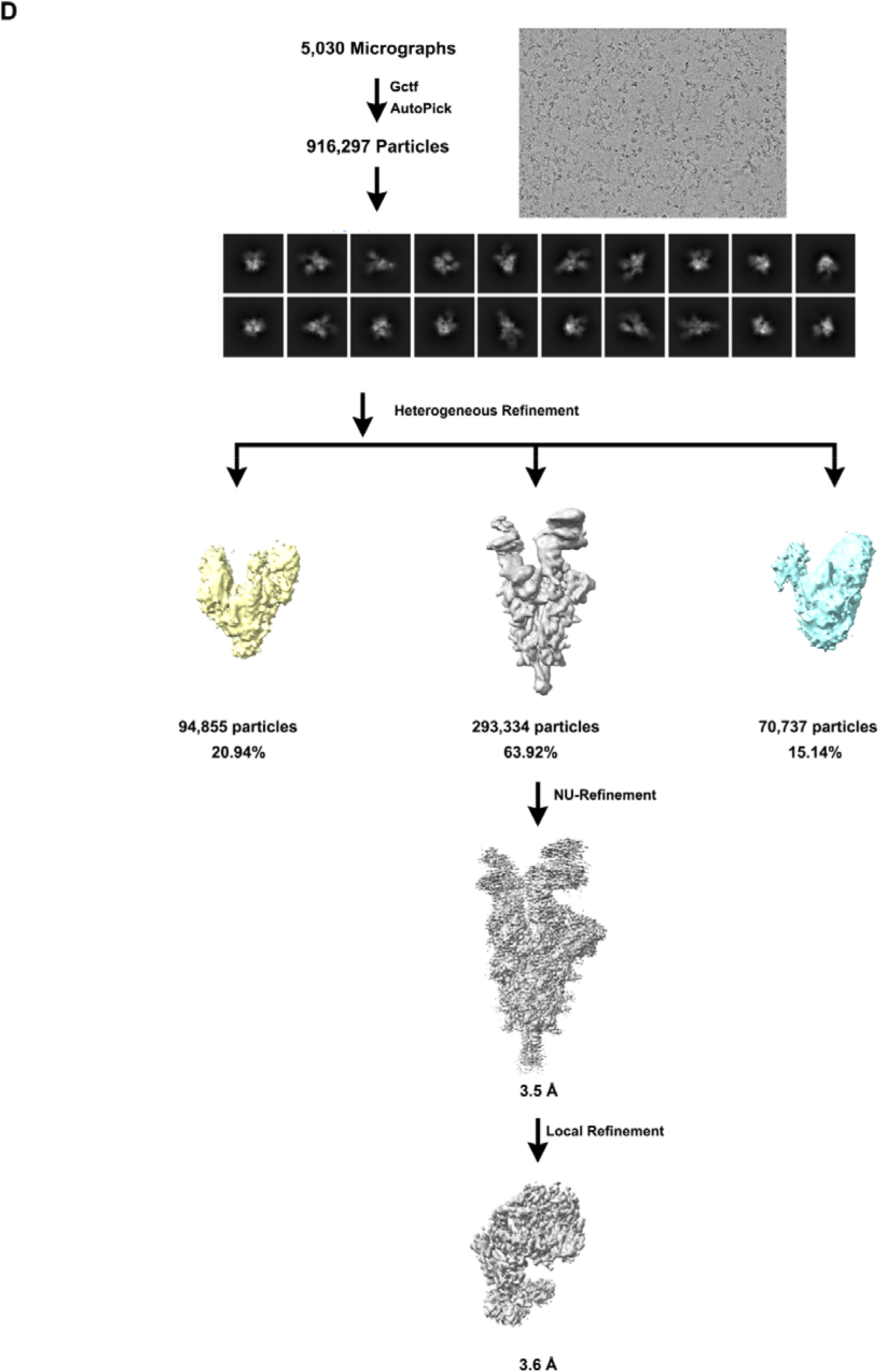
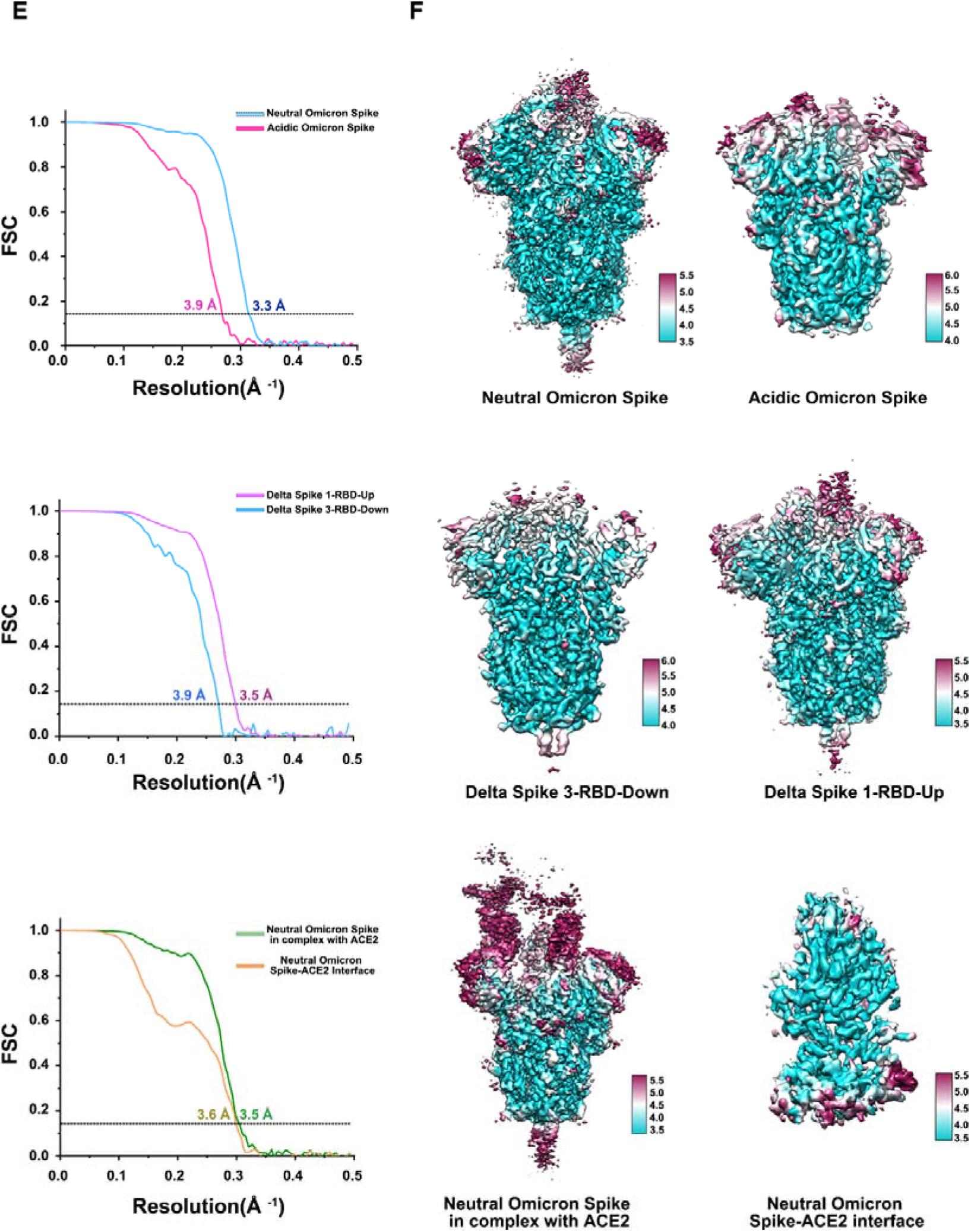
Flowchart for Cryo-EM data processing and resolution estimation of the EM maps. Flowcharts for (**A**) neutral Omicron Spike protein, (**B**) acidic Omicron Spike protein, (**C**) Delta Spike protein, (**D**) neutral Omicron Spike protein in complex with hACE2. (**E**) The gold-standard FSC curves of overall maps of neutral Omicron Spike protein, acidic Omicron Spike protein, Delta Spike protein, neutral Omicron Spike protein in complex with hACE2 and local maps of interfaces. (**F**) Local resolution assessments of cryo-EM maps using ResMap are shown.

**Figure S3.**
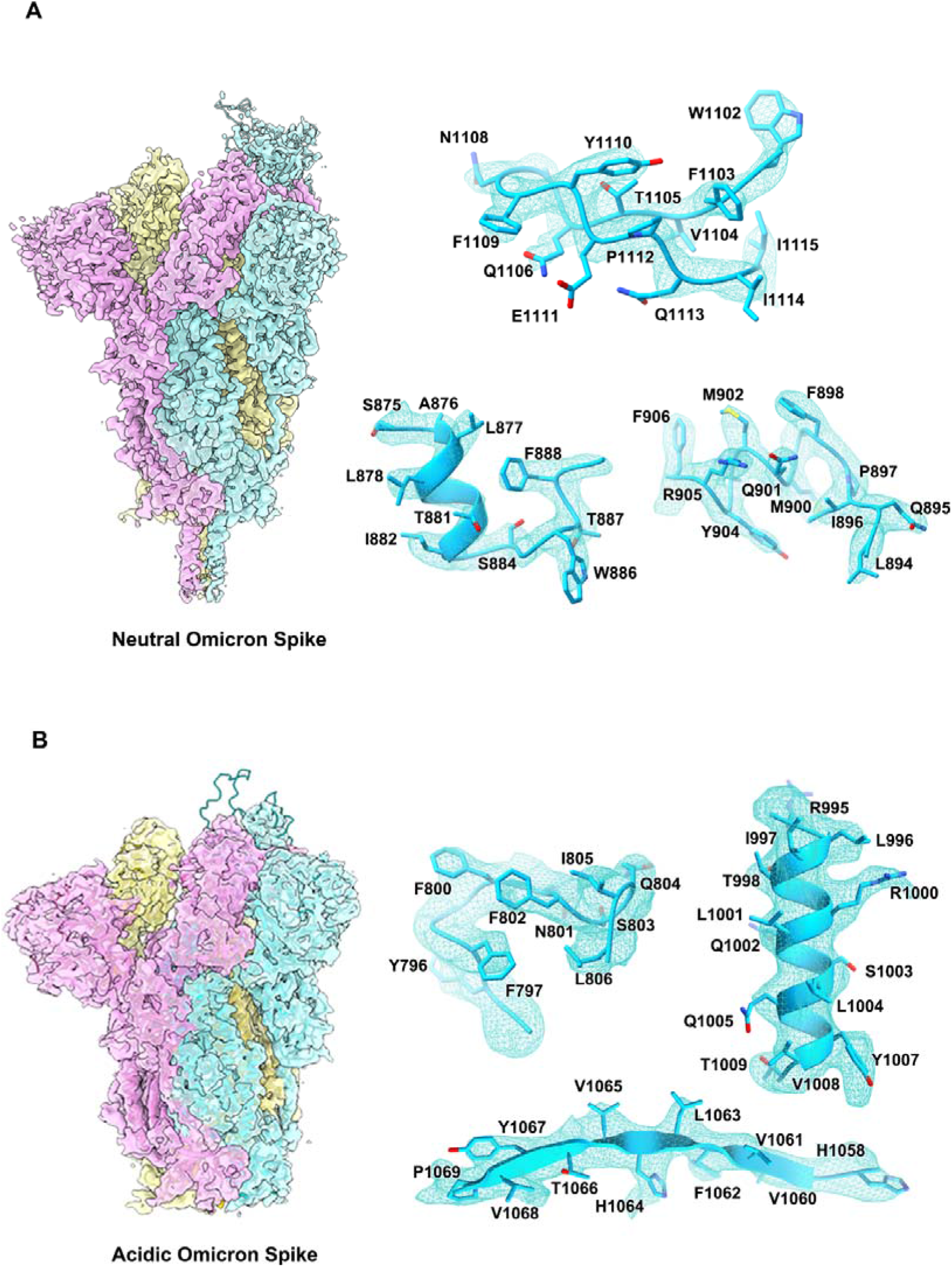
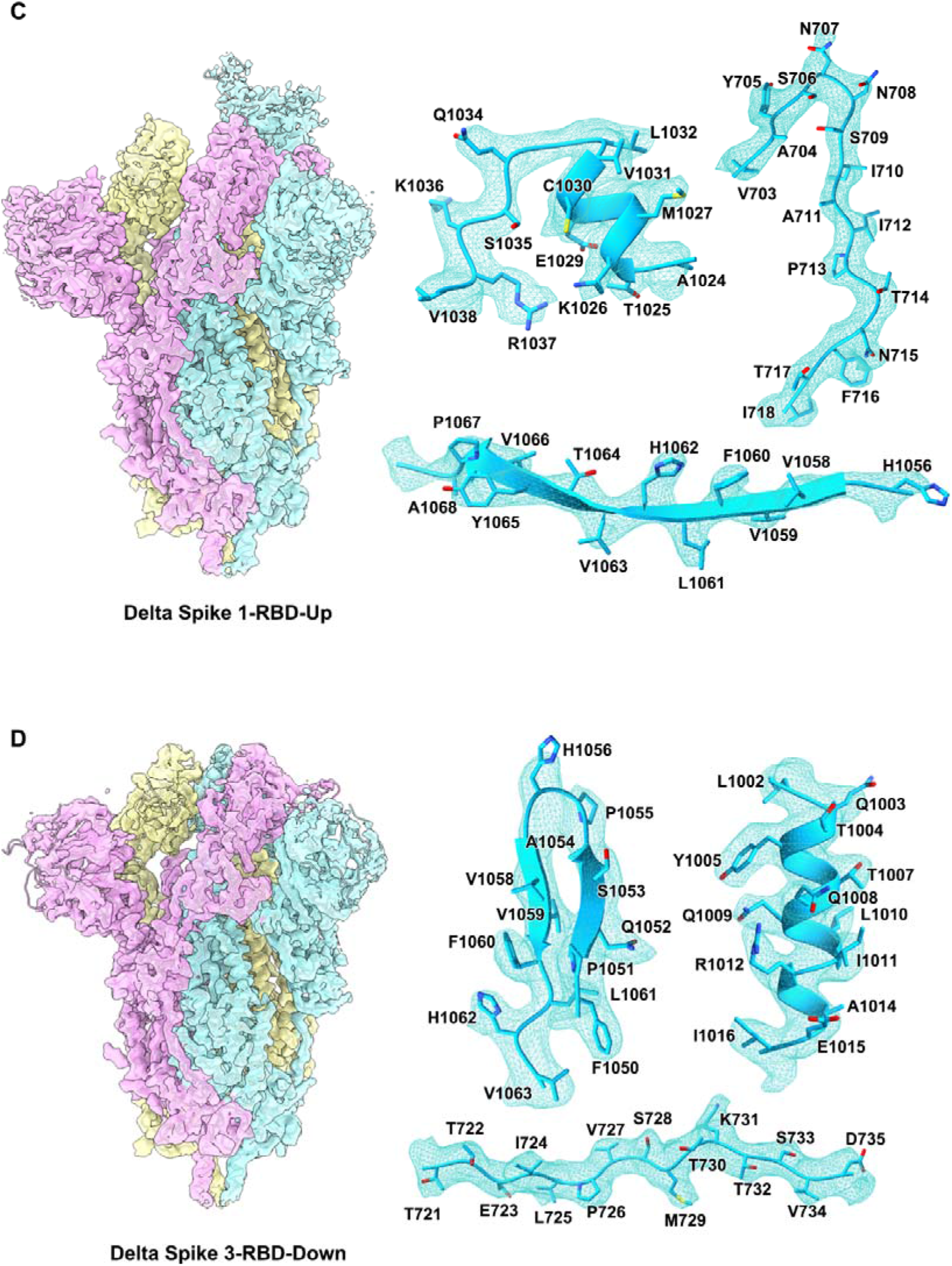
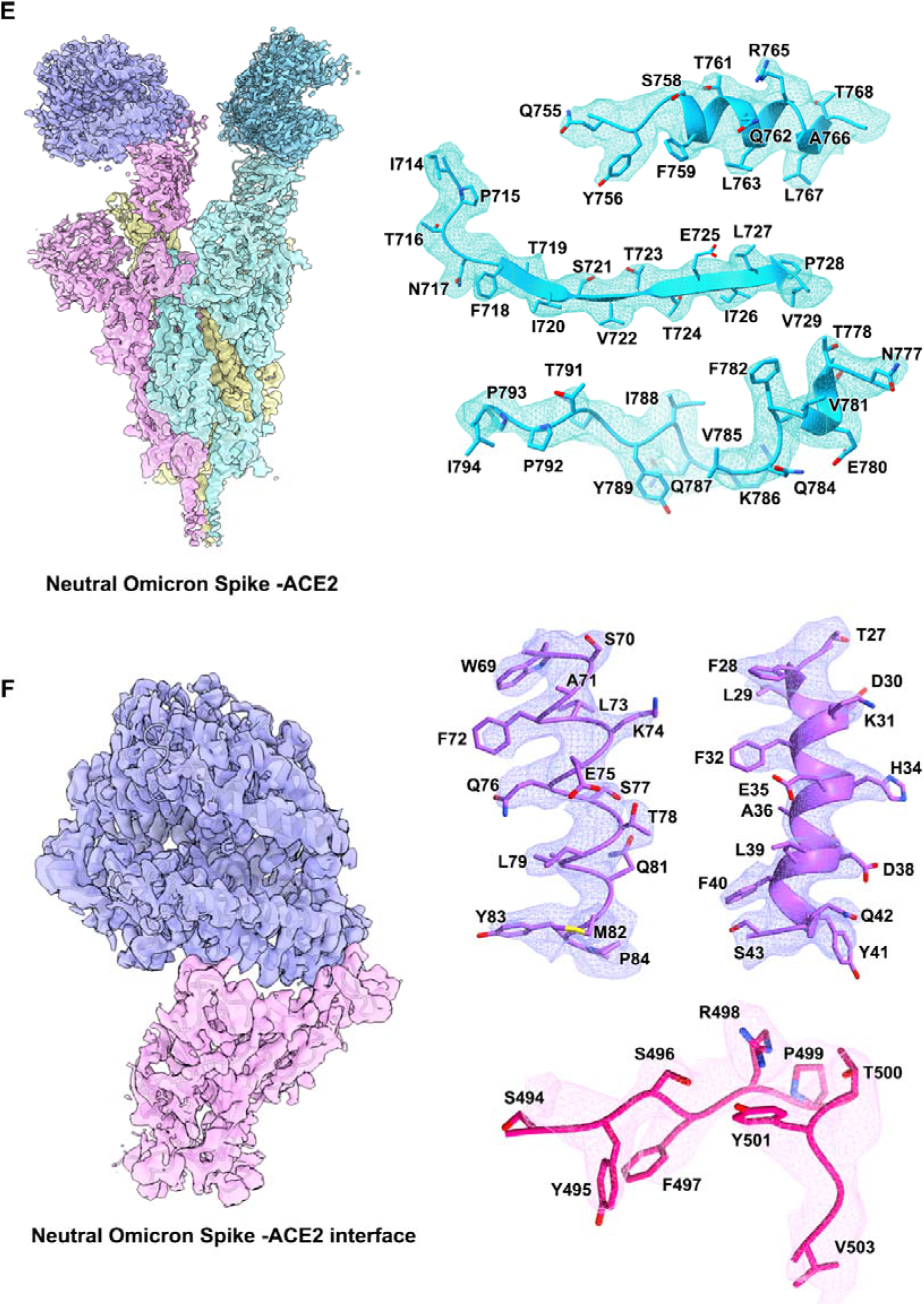
Density maps and atomic models. Cryo-EM maps of (**A**) neutral Omicron Spike protein, (**B**) acidic Omicron Spike protein, (**C**) Delta Spike protein 1-RBD-up, (**D**) Delta Spike protein 3-RBD-down, (**E**) neutral Omicron Spike protein in complex with hACE2 and (**F**) local maps of interfaces. Residues are shown as sticks with oxygen colored in red, nitrogen colored in blue and sulfurs colored in yellow. Three different protomers of S trimer are colored in cyan, pink and yellow, respectively.

**Figure S4.**
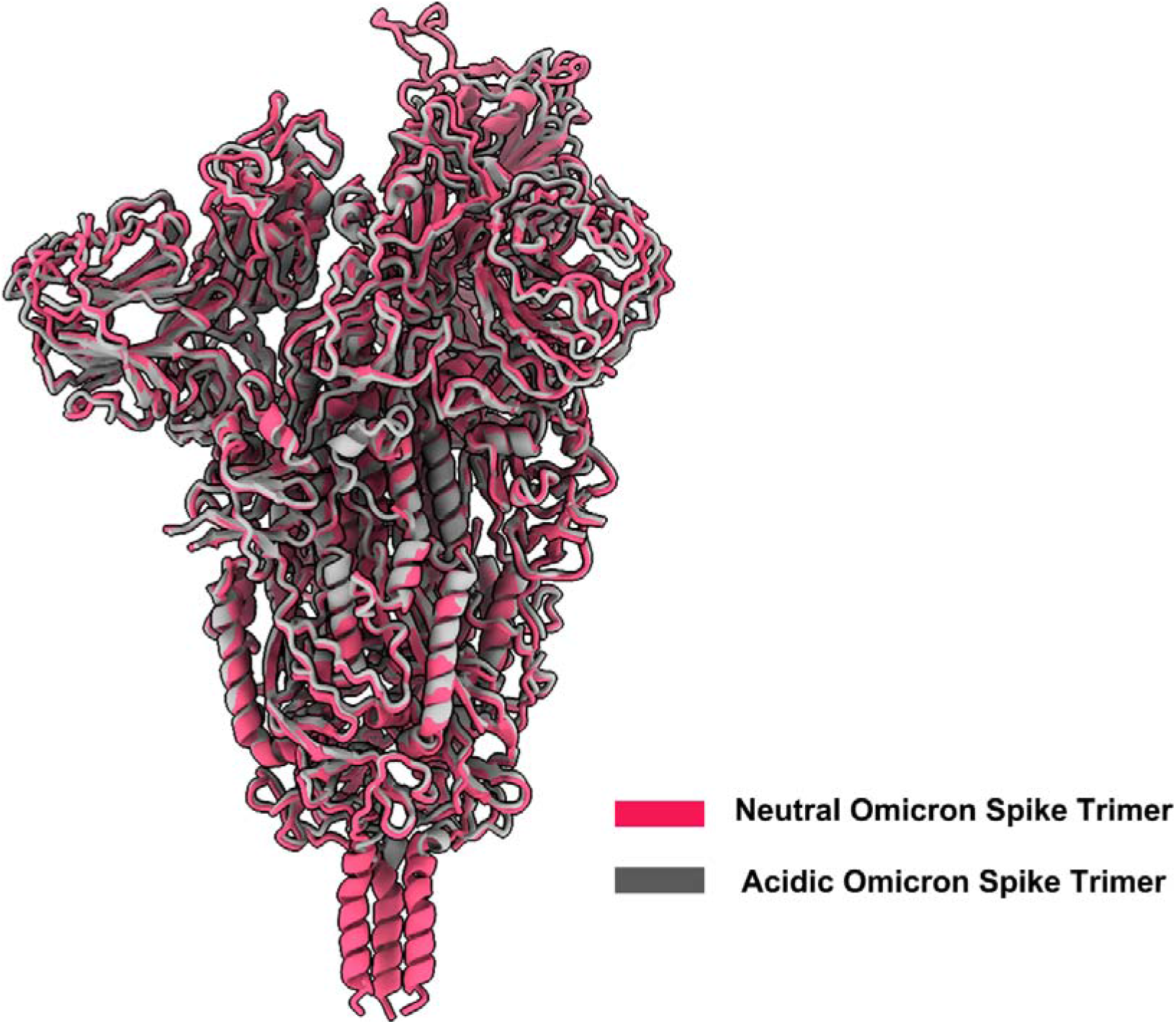
Superimposition of overall structures of neutral and acid Omicron S-trimers. The structure of acid Omicron S-trimer (grey) is superimposed to neutral Omicron S-trimer (red) and no features except some disorder on NTD and RBM is observed.

**Figure S5.**
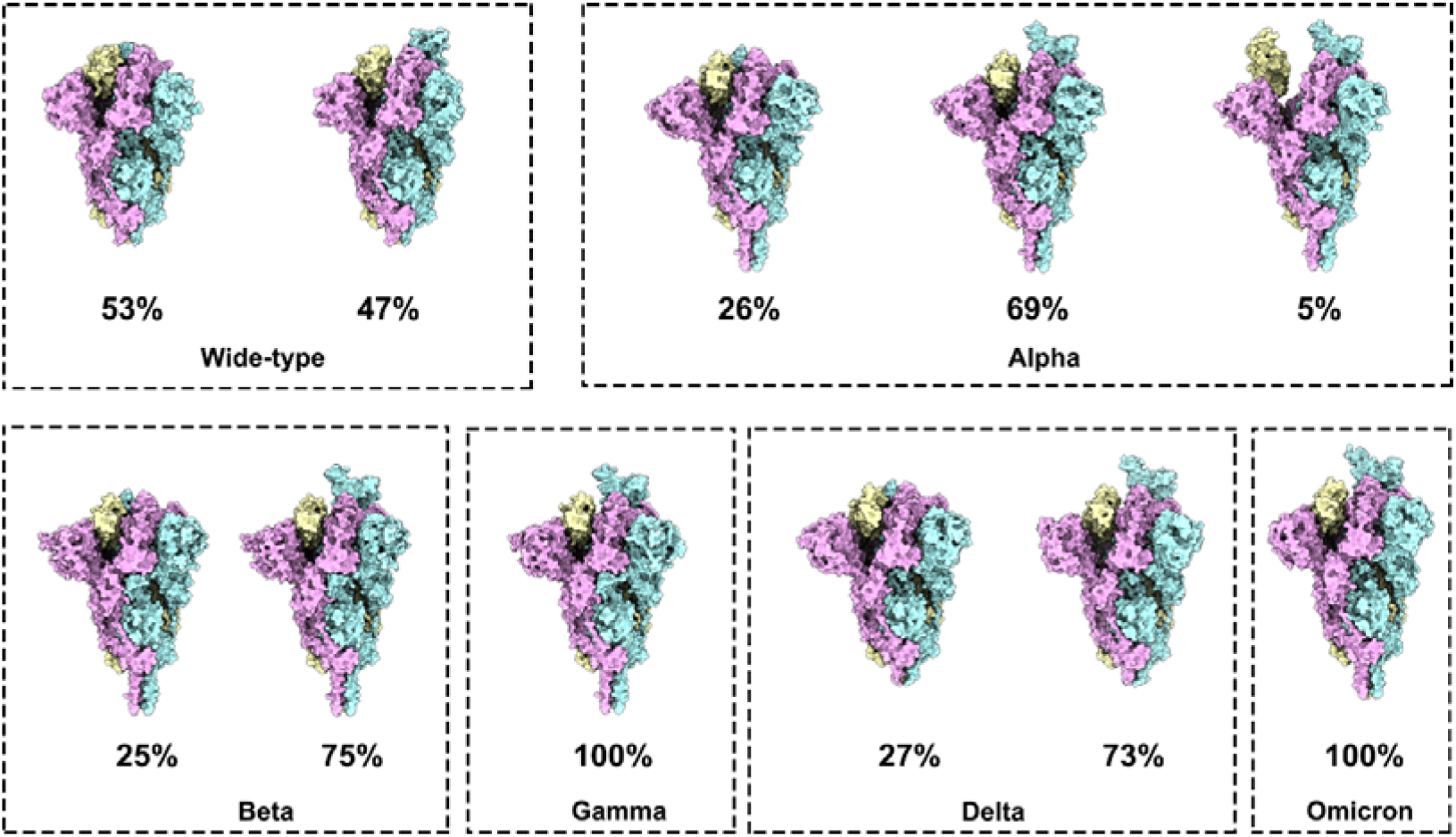
Conformational states of Wild-type and variant S trimer. Surface of Wild-type S trimer, Alpha S trimer, Beta S trimer, Gamma S trimer, Delta S trimer and Omicron S trimer shown with each protomer in different color (cyan, pink and yellow). The ratio of S protein in different conformations is shown in the figure.

**Figure S6.**
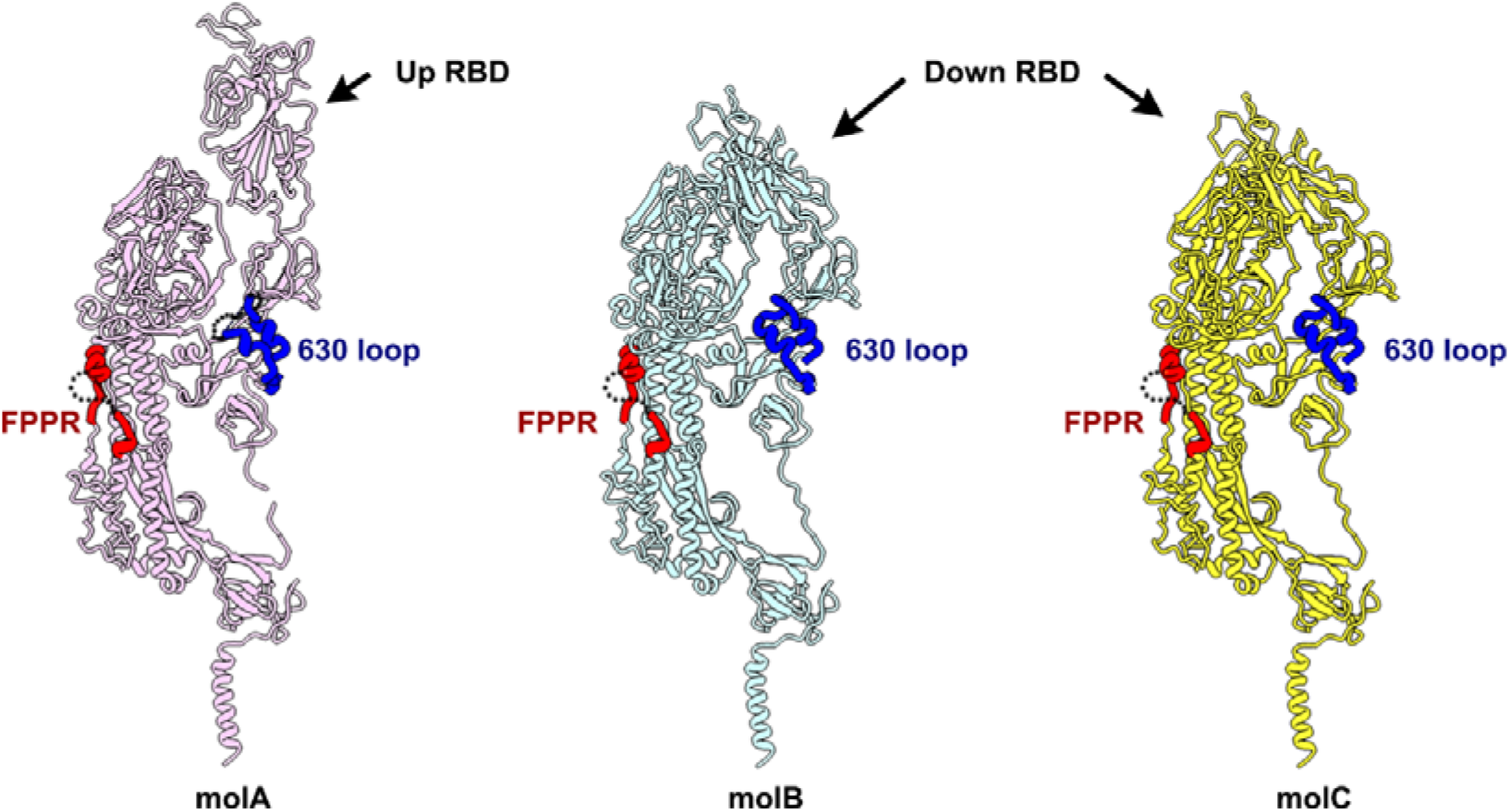
Structures of 630 loop and FPPR in Omicron S trimer. Three protomers (molA, magenta; molB, cyan and molC, yellow) of Omicron S trimer are shown as ribbon in the same orientation. Structures of FPPR and 630 loop on each protomer are accentuated in bold red and bold blue, respectively.

**Figure S7.**
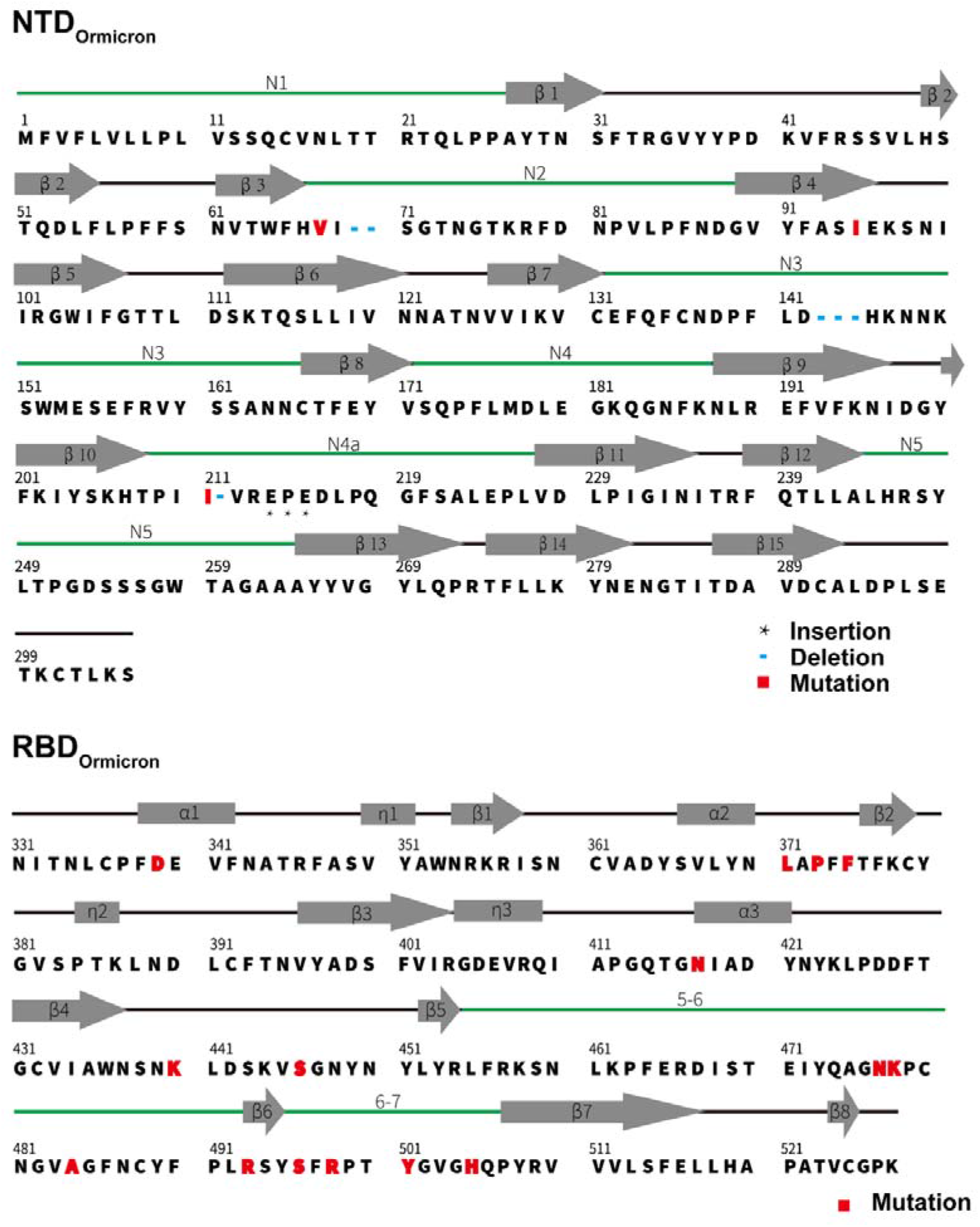
Secondary structure of the Omicron RBD and NTD. Primary sequence and secondary structural elements of the Omicron RBD and NTD, the five-pointed star represents the inserted residues, the blue short line represents the deleted residues and the red box represents the mutated residues.

**Figure S8.**
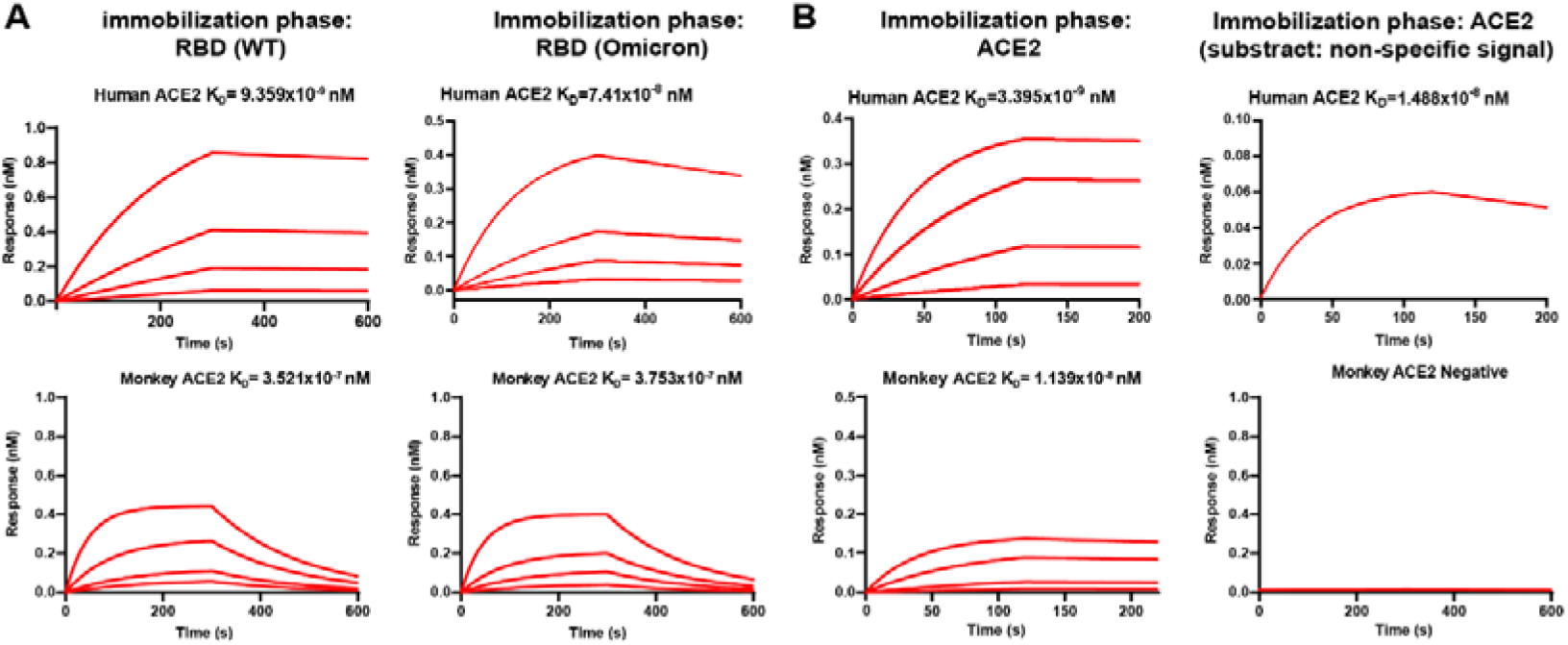
Affinity of Omicron RBD to ACE2 from different species. (A) The wild-type RBD (left) and omicron RBD (right) were loaded onto the biosensor, the human or monkey ACE2 containing buffer solution was passed over the bound RBD. (B) The human or monkey ACE2 was immobilized onto biosensor, while the Omicron RBD was used as the analyte (left). The final corrected curves after removing non-specific bindings are shown (right).

**Figure S9.**
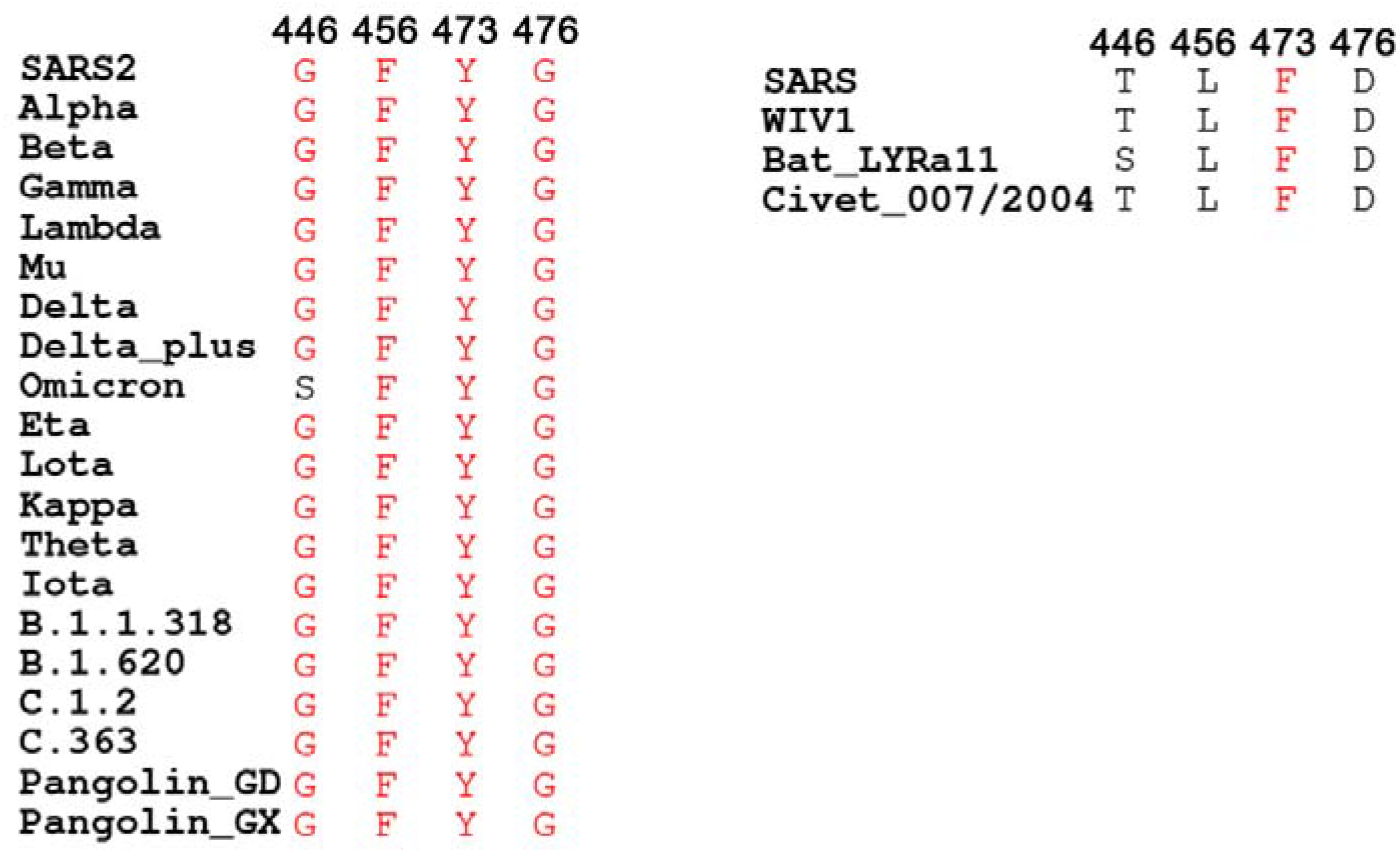
Multiple sequence alignment of residues involved in ACE2 binding from 25 representative sarbecovirus members.

**Figure S10.**
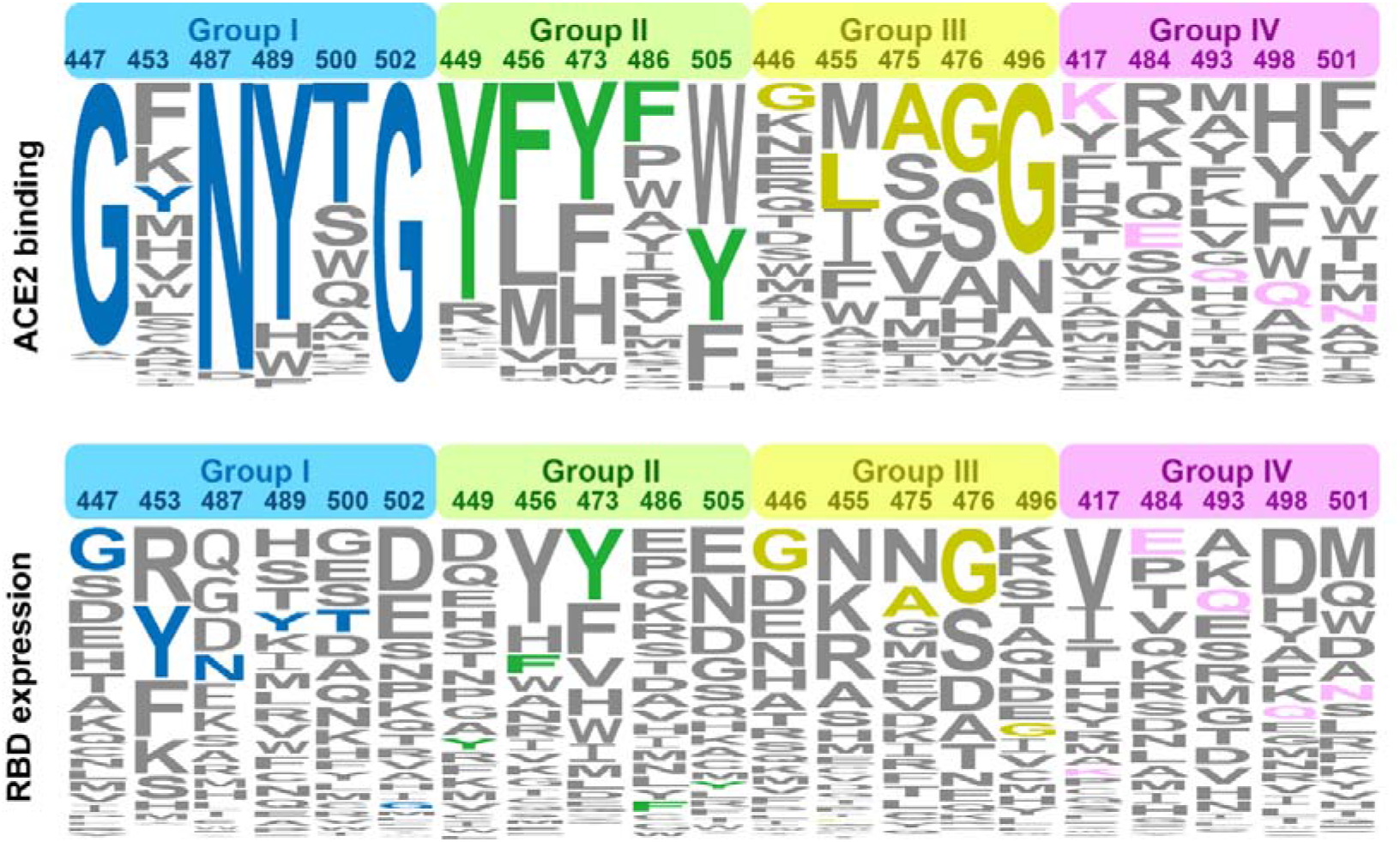
Modified Logo Plot from deep mutagenesis scanning datasets Represents the effects of Mutations present in key residues on ACE2 binding and RBD expression (37). The height of an amino acid code indicates its preference on a given site of RBM with respect to ACE2 binding (top) and RBD expression (bottom), as is shown previously. Residues are classified into four groups, ie., the identical residues, homologous residues, conditionally altered residues and diverse residues, which are colored in blue, green, orange and red, respectively.

## Supplemental Table

**Table S1.** Cryo-EM data collection and atomic model refinement statistics.

| Name | Neutral<br>Omicron<br>Spike | Acidic<br>Omicron<br>Spike | Delta Spike<br>(1 RBD Up) | Delta Spike<br>(3 RBD Down) | Neutral<br>Omicron<br>Spike in<br>complex<br>with<br>ACE2 | Neutral<br>Omicron<br>Spike-ACE2<br>interface |
| --- | --- | --- | --- | --- | --- | --- |
| **Data collection** |  |  |  |  |  |  |
| Microscope | FEI<br>Titan<br>Krios | FEI<br>Titan<br>Krios | FEI Titan<br>Krios | FEI Titan Krios | FEI Titan<br>Krios | FEI Titan<br>Krios |
| Camera | K3 | K3 | K3 | K3 | K3 | K3 |
| Voltage (kV) | 300 | 300 | 300 | 300 | 300 | 300 |
| Total dose (e<sup>-</sup><br>/Å<sup>2</sup>) | 60 | 60 | 60 | 60 | 60 | 60 |
| Symmetry<br>imposed | C1 | C1 | C1 | C3 | C1 | C1 |
| **Data Processing** |  |  |  |  |  |  |
| Micrographs<br>(Total) | 1,777 | 6,050 | 3,106 | 3,106 | 5,030 | 5,030 |
| Micrographs<br>(Used) | 1,777 | 6,050 | 3,106 | 3,106 | 5,030 | 5,030 |
| Particles<br>selected | 830,943 | 1,789,492 | 1,317,048 | 1,317,048 | 916,297 | 916,297 |
| Particles<br>included in<br>final<br>reconstruction | 221,744 | 371,808 | 491,247 | 171,670 | 293,334 | 293,334 |
| **Reconstruction** |  |  |  |  |  |  |
| Pixel size (Å) | 1.07 | 1.07 | 1.07 | 1.07 | 1.07 | 1.07 |
| Defocus<br>range (µm) | -1.5--2.5 | -1.5--2.5 | -1.5--2.5 | -1.5--2.5 | -1.5--2.5 | -1.5--2.5 |
| Resolution<br>(Å) (FSC =<br>0.143) | 3.3 | 3.9 | 3.5 | 3.9 | 3.5 | 3.6 |
| **Model Refinement** |  |  |  |  |  |  |
| Clashscore | 7.1 | 12.27 | 7.83 | 7.93 | 12.9 | 3.68 |
| Rotamer<br>outliers (%) | 0.00 | 0.03 | 0.00 | 0.04 | 0.05 | 0 |
| Molprobity<br>score | 1.77 | 2.1 | 1.79 | 1.81 | 2.09 | 1.49 |
| **Ramachandran statistics (%)** |  |  |  |  |  |  |
| Most favored<br>(%) | 94.43 | 91.69 | 94.67 | 94.54 | 92.58 | 95.18 |
| Allowed (%) | 5.51 | 8.28 | 5.33 | 5.46 | 7.33 | 4.82 |
| Outliers (%) | 0.06 | 0.03 | 0 | 0 | 0.09 | 0 |
| **R.m.s.deviations** |  |  |  |  |  |  |
| Bond lengths<br>(Å) | 0.003 | 0.003 | 0.003 | 0.003 | 0.009 | 0.006 |
| Bond angles<br>(°) | 0.56 | 0.611 | 0.568 | 0.562 | 0.641 | 1.155 |

